# Branch Length Transforms using Optimal Tree Metric Matching

**DOI:** 10.1101/2023.11.13.566962

**Authors:** Shayesteh Arasti, Puoya Tabaghi, Yasamin Tabatabaee, Alan K. Mayer, Siavash Mirarab

## Abstract

The abundant discordance between evolutionary relationships across the genome has rekindled interest in methods for comparing and averaging trees on a shared leaf set. However, compared to tree topology, where much progress has been made, handling branch lengths has been more challenging. Species tree branch lengths can be measured in various units, often different from gene trees. Moreover, rates of evolution change across the genome, the species tree, and specific branches of gene trees. These factors compound the stochasticity of coalescence times and estimation noise, making branch lengths highly heterogeneous across the genome. For many downstream applications in phylogenomic analyses, branch lengths are as important as the topology, and yet, existing tools to compare and combine weighted trees are limited. In this paper, we address the question of matching one tree to another, accounting for their branch lengths. We define a series of computational problems called Topology-Constrained Metric Matching (TCMM) that seek to transform the branch lengths of a query tree based on a reference tree. We show that TCMM problems can be solved efficiently using a linear algebraic formulation coupled with dynamic programming preprocessing. While many applications can be imagined for this framework, we explore two applications in this paper: embedding leaves of gene trees in Euclidean space to find outliers potentially indicative of errors, and summarizing gene tree branch lengths onto the species tree. In these applications, our method, when paired with existing methods, increases their accuracy at limited computational expense.

## Introduction

Discordance among evolutionary histories across the genome (Maddison, 1997) is now understood to be pervasive and has motivated the study of region-specific histories (gene trees) and the speciation history (the species tree). Having alternative trees leads to a need for comparing and combining trees. We need metrics to quantify tree dissimilarity, algorithms for mapping gene trees to the species tree to reconcile them (Doyon et al., 2011), and median tree methods to combine gene trees into a single species tree (Barthélemy and McMorris, 1986). Solutions to these problems have been developed and are widely adopted. Measures of tree difference such as Robinson and Foulds (1981) (RF), quartet distance (Estabrook et al., 1985), and matching split (MS) (Bogdanowicz and Giaro, 2012) are commonly used to quantify gene tree differences. Biologists often use reconciliation methods in comparative genomics to understand the evolution of gene function. Inferring the species tree by finding the tree with the minimum total quartet distance to gene trees (quartet median tree) has been widely adopted due to its statistical consistency under several models of gene tree discordance (Mirarab et al., 2021).

The rich literature on comparing gene trees and species trees has focused on tree topology far more than branch lengths. Discordance, however, is not just limited to topology. One of the main causes of discordance is incomplete lineage sorting (ILS) as modeled by the multi-species coalescent (MSC) model (Degnan and Rosenberg, 2009). Under MSC, internal nodes of a gene tree correspond to coalescence events that predate species divergences. As a result, gene tree branch lengths are guaranteed to differ from the species tree even if there is no topological discordance (a phenomenon Edwards and Beerli (2000) referred to as species versus genic divergences). Beyond true biological differences, challenges in inferring gene trees compound the problem. From sequences alone, we can only measure branch lengths in the unit of the expected number of substitutions per site (SU), which are a function of not just divergence times but also substitution rates, which vary across the tree, across the genome, or both (Kumar, 2005; Lopez et al., 2002; Rasmussen and Kellis, 2007). Furthermore, gene trees are often inferred from short sequences (to avoid recombination within a gene), which makes gene tree estimates noisy, especially for short branches.

Branch lengths are as important as topology to many downstream applications such as dating, diversification studies, comparative genomics, trait evolution, and even microbiome analyses. Nevertheless, the myriad sources of heterogeneity (ILS, rate change, noise) have made it challenging to reconcile gene tree branch lengths with each other and with the species tree. Bayesian co-estimation methods (e.g., Boussau et al., 2013; Heled and Drummond, 2010; Liu, 2008) account for branch lengths by jointly modeling gene trees and species trees and introducing relevant parameters. These methods tend to be accurate (Yang and Warnow, 2011) and readily produce a reconciliation of species trees and gene trees. However, they do not scale to large datasets (Ogilvie et al., 2017; Zimmermann et al., 2014). Moreover, since complex rate models further reduce scalability, these methods often model only limited rate heterogeneity.

The alternative two-step approach of inferring gene trees and then summarizing them to find the species tree often ignores branch lengths. Theoretically justifiable methods that consider branch lengths in inferring species trees (e.g., GLASS and STEM; Kubatko et al., 2009; Mossel and Roch, 2010) have been less accurate than topology-focused methods given estimated gene trees (DeGiorgio and Degnan, 2014; Yang and Warnow, 2011), partially due to challenges mentioned earlier (Huang et al., 2010). Others have focused on gene tree branch lengths (SU) for rooting them (Mai et al., 2017; Tria et al., 2017), for finding outliers indicative of errors (de Vienne et al., 2012; Mai and Mirarab, 2018), for clustering gene trees (Gori et al., 2016), and for summarizing them to infer the species tree branch lengths (Binet et al., 2016; Tabatabaee et al., 2023). These attempts have been more successful but lack an underlying mechanism for matching trees that can be used across applications. Each deals in some way with the underlying difficulty that two trees that differ in both topology and branch length are hard to compare since the same branches are not present in both (see Kuhner and Yamato (2015) for tree comparison metrics that include branch length).

To arrive at a general method for comparing trees while accounting for branch lengths, we propose a framework that decouples topological changes and branch length changes to an adjustable extent. We use a linear algebraic formulation based on patristic distances (pairwise path lengths) to define an optimization problem called Topology-Constrained Metric Matching (TCMM), which is inspired by the least-square distance-based methods (Cavalli-Sforza and Edwards, 1967; Rzhetsky and Nei, 1991, 1992, 1993). The goal of TCMM is to assign branch lengths to a query tree such that its patristic distances match a set of one or more reference trees as much as possible without changing its topology. Using an additional regularization term, we can balance the competing goals of matching the reference tree and keeping the original branch length patterns of the query tree. All TCMM problems are convex, require preprocessing doable in quadratic time and memory, and are solvable using standard gradient descent techniques. While this ability to align the metric structure of one tree with another offers a powerful tool, in this paper, we focus on two particular applications: 1) embedding gene tree leaves in a shared space to enhance outlier detection, and 2) transferring gene tree branch lengths to a species tree. We show that in both applications, using TCMM to match gene tree branch lengths, when paired with existing methods, can improve their accuracy.

## Material and Methods

### Problem formulation

For an unrooted tree *T* = (*V*_*T*_, *E*_*T*_), we let *L*_*T*_ denote the leaf set and *P*_*T*_ (*v, v*^′^) denote the path between any pairs of vertices *v, v*^′^ ∈ *V*_*T*_. For a weighted tree, each edge *e* ∈ *E*_*T*_ has length *w*_*T*_ (*e*) ∈ ℝ^+^. We use *T* (*T*) to denote the set of all weighted trees with the same topology as tree *T*. A weighted tree *T* with *N* vertices indexed by [*N*] = {1,…,*N*} defines a metric *d*_*T*_ : *V*_*T*_ *× V*_*T*_ → ℝ^+^ where 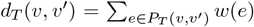. We define the *weight* vector 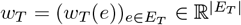 as a vector of all the weights indexed by edges and, with a slight abuse of notation, the *pairwise distance* vector 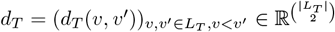 as a vector of all patristic distances indexed by pairs of nodes. We define a binary *path matrix A*_*T*_ of size 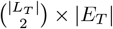 where rows correspond to pairs of vertices in *L*_*T*_ and columns represent the edge set. For two distinct vertices *v, v*^′^ ∈ *L*_*T*_ and edge *e* ∈ *E*_*T*_, an element ((*v, v*^′^), *e*) of *A*_*T*_ is one if and only if the edge *e* is on the path between *v* and *v*. Otherwise, it is equal to zero. The path matrix *A*_*T*_ completely encodes the topology of *T*.

Consider a query tree *T* and a reference weighted tree *R*, both on a shared leafset *L*. For any tree *T* ^′^ with the same topology as *T*, let cost(*T*; *R*) measure the branch length discrepancy of *T* to the reference tree *R*. For any such cost function, we can seek 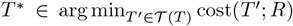. Since the topology is fixed in this problem, we can reparameterize the cost with respect to the weight vector *w*_*T*_ to get 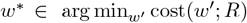, where 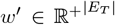 are constrained to be nonnegative. In this paper, we focus on a particular cost function to quantify the discrepancy of metric information between the two trees and successively refine it. Akin to least squares error distance-based tree estimation, we set

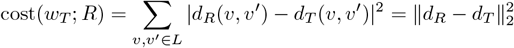

as the square of the *f*_2_ distance between pairwise distance vectors. As it has long been appreciated (Cavalli-Sforza and Edwards, 1967) (see also Desper and Gascuel, 2003), we can find a separable form for pairwise distances in terms of topology and edge weights. For *v, v* ^′^ ∈ *L*, we can write 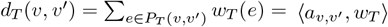 where 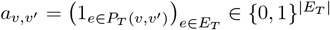 and 1_*x*_ is the indicator function. Note that 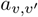 is a row of the path matrix *A*_*T*_. Thus, for any tree *T* = (*V*_*T*_, *E*_*T*_), we have *d*_*T*_ = *A*_*T*_ *w*_*T*_, and

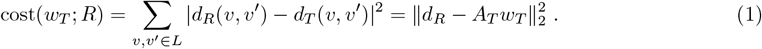

We can now define an ordinary least squares (OLS) problem similar to Cavalli-Sforza and Edwards (1967) (and formulated more clearly by Rzhetsky and Nei (1991)):

**Problem 1** (Topology-Constrained Metric Matching— under 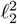). *Let T, R be weighted trees with edge weights w*_*T*_ *and w*_*R*_ *on the common leafset L with at least two elements. The weights of the closest tree to R matching T in topology with edge weights lower bounded by ε* ≥ 0 *is:*

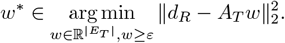

The cost function in Problem 1 is agnostic to the original branch lengths of *T, w*_*T*_, and thus, it can dramatically alter them. For many applications, it is desirable to retain the original branch lengths to some user-adjustable extent. We can accommodate that goal by adding a regularizing term. For an optimal edge weight vector 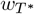, we define the *adjustment rate vector* by element-wise dividing 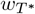 and *w*_*T*_, to get 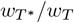. To incorporate *w*_*T*_ in the problem statement, we penalize the deviation of estimated 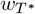 from the initial weight vector *w*_*T*_, save for a total scaling factor shared by all branches, as in the following problem.

**Problem 2** (Regularized Topology-Constrained Metric Matching — under 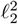). *Let T, R be weighted trees with edge weight functions w*_*T*_ *and w*_*R*_ *on the common leafset L. Let λ be a fixed regularization coefficient and σ*^2^(*w/w*_*T*_) *denote the variance of the adjustment rates vector w/w*_*T*_. *We seek the tree T* ^*^ ∈ *T* (*T*) *with edge weights*

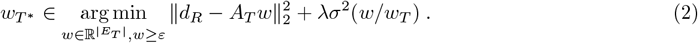

The regularizing term, which is a quadratic function of the weight vector *w*, leads to lowered variance in the adjustment rate vector, *w/w*_*T*_, and thus, lowered change in *relative* branch lengths compared to *T*. Clearly, if the variance of adjustment rate values is 0, the tree *T* has been simply scaled up or down, without any real change to its relative branch lengths. Higher *λ* values encourage less change to the relative branch lengths of *T*. As *λ* → ∞, all rate multipliers converge to a solution that simply scales branch lengths of *T* with a fixed global rate *r*\*:

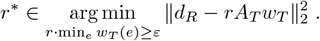

To make *λ* values more interpretable across all datasets with varying sizes and heights, we first compute the optimal branch lengths *w*^*^ according to Problem 1. We then reformulate Problem 2 as:

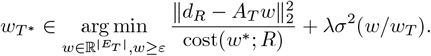

where cost(*w*^*^; *R*) is defined in Eq. (1). This adjustment ensures that the first term is scale-free and provides an initial solution that can be used as input for the optimizer. We will use this revised formulation of the Regularized TCMM Problem in all experiments. We can also extend to multiple inputs.

**Problem 3** (Topology-Constrained Consensus Metric Matching). *Let* {*R*_*k*_ : *k* ∈ [*K*]} *be a set of weighted reference trees, each with weights w*_*k*_. *Given a tree T, we seek to find the optimal (adjusted) tree T* ^*^ *with the same topology as T and weights* 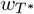 *such that:*

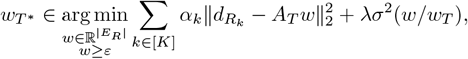

*where α*_*k*_ *is a nonnegative weight controlling the contribution of reference tree R*_*k*_ *in the cost function, λ>* 0 *is a regularizing coefficient, σ*^2^(*w/w*_*T*_) *penalizes the weights that result in a large variance for the adjustment rate vector w/w*_*T*_, *and ε* ≥ 0 *is a lower bound for the estimated edge weights*.

Remark 1 (supplementary material) shows that problem 3 can be solved by solving Problem 2 after averaging distances across *k* genes.

### A Scalable Convex Optimization Solution

#### TCMM problems are all convex

We can rewrite the cost functions in Problems 1–3 in quadratic form. For simplicity, we use *A* = *A*_*T*_ and *d* = *d*_*R*_. Note that Problem 1 minimizes 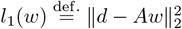. We convert *l*_1_ to the quadratic form

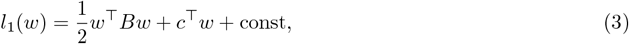

where *B* = 2*A*^*T*^*A* is a positive semidefinite matrix, *c* = −2*A*^*T*^*d*, and const 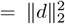. Since *B* ⪰ 0 by construction, *l*_1_ is a convex function of the weight parameter *w*. Adding the regularization term *σ*^2^(*r*) — where *r* = *w/w*_*T*_ — does not change the convexity, as we can convert it to the following quadratic form:

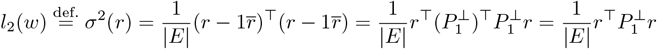

where 1 ∈ ℝ^|*E*|^ is the vector of all ones, *r* is a scalar that is the average value of elements in 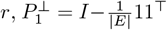. It is a known fact that 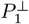 is an orthogonal projection operator, that is, 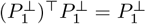. Furthermore, it is a positive semidefinite matrix with 0 − 1 eigenvalues. Therefore, the regularizing term *σ*^2^(*w/w*_*T*_) is indeed a convex function of the weight parameter *w*. Thus, Problems 1,2, and 3 all admit globally optimal solutions.

**Proposition 1**. *The cost functions in Problems 1, 2, and 3 are convex functions of w defined over nonempty, closed, and convex constraint sets. Hence, each optimization problem admits a global minimum (though not necessarily a unique minimizer), and the projected gradient descent method converges to such a solution*.

Proofs are provided in the supplementary material, Appendix A.5. By Proposition 1, we can use projected gradient descent to solve Problems 1 – 3 using the gradient terms shown in the proof. We implemented and optimized the quadratic functions *l*_1_ and *l*_2_ using built-in functions in the Python package `CVXPY`. Our empirical results suggest a sub-cubic running time (≈ *O*(*N* ^2.74^)) for this implementation (Fig. S1a).

#### Dynamic programming enables calculating *B* and *c* in quadratic time

Our formulation does not need to ever compute *A*_*T*_ and *d*_*R*_ — whose dimensions grow cubically with *N*. Rather, it needs the matrix *B*, which grows quadratically, and the vector *c*, which grows linearly with *N*. Naively computing 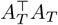 from its building blocks requires *O*(*N* ^4^) time and *O*(*N* ^3^) memory. Similarly, computing 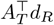 naively requires *O*(*N* ^3^) time and *O*(*N* ^3^) memory. However, we can compute *B* in quadratic, and *c* in sub-quadratic time and memory using dynamic programming without ever computing *A*_*T*_ and *d*_*R*_.

#### Computing 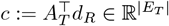

Bryant and Waddell (1998) propose a quadratic algorithm to compute 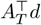 for any arbitrary dissimilarity matrix *d*. However, in our case, since *d*_*R*_ comes from a tree, we can design a faster algorithm to compute 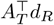. Each edge *e* ∈ *E*_*T*_ corresponds to a bipartition of leaves of *T*. For an arbitrary rooting of *T*, we define *C*_*T*_ (*e*) as the set of leaves of *T* below *e*. Let *e*^′^ and *e*^″^ be the children of *e*. As Bryant and Waddell (1998) noted, the element of *c* that corresponds to *e* is

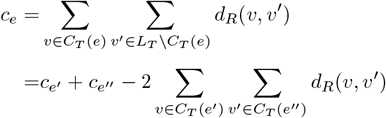

where the second equation gives a recursive algorithm. For an arbitrary distance matrix, the double-sum in the recursion requires *O*(*N* ^2^) over the entire tree. While *O*(*N* ^2^) will be dominated by the optimization step, we note that in Problem 3, that calculation of *c* will have to be repeated *k* times, which means without further improvements, calculation of *c* could become dominant for large *k*. Luckily, when *d*_*R*_ is defined on a given *R*, we can do better. We have designed an algorithm (see Algorithm S1) to compute that term using |*C*_*T*_ (*e*^′^) + *C*_*T*_ (*e*^″^)| leaf-to-root traversals on *R*. Each traversal takes *O*(*H*_*R*_), where *H*_*R*_ is the height of *R*, and using the recursive equation above, we need exactly *N* · *H*_*T*_ such traversals, where *H*_*T*_ is the height of *T*. Therefore, the total running time of this algorithm is *O*(*N* · *H*_*T*_ · *H*_*R*_). While this algorithm can be as slow as *O*(*N* ^3^) for unbalanced trees, it would require *O*(*N* log^2^(*N*)) for sufficiently balanced trees, often encountered in practice. Our empirical results (Fig. S1b) show that on average it is sub-quadratic (close to *O*(*N* ^1.4^)). Note also that for unbalanced input (defined as *H*_*T*_ · *H*_*R*_ *> N*), we switch to the quadratic algorithm of Bryant and Waddell (1998), thus ensuring the running time is never worse than quadratic.

In the consensus Problem 3, if no distances are missing, we can simply average the terms 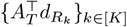 instead of averaging distance vectors 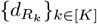. The consensus cost function allows this:

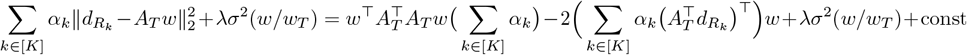

#### Computing 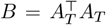

Given the definition of the path matrix *A*_*T*_, it can be easily deduced that for two edges *e* and 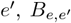 is the number of pairs of leaves *v, v*^′^ ∈ *L* where *e, e*^′^ ∈ *P*_*T*_ (*v, v*^′^). We easily pre-compute |*C*_*T*_ (*e*)| for each edge *e* ∈ *E*_*T*_ in linear time and the least common ancestor (LCA) of all pairs of nodes of *T* in quadratic time. With these, we can easily compute 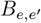 for all *e, e*^′^ ∈ *E*_*T*_ in quadratic time using simple calculations given in Algorithm S1 (supplementary material), as confirmed by our empirical analysis (Fig. S1c).

We have assumed so far that query and reference trees have the same species, but this will not always be the case. Gene trees almost always miss some taxa due to gene loss or data incompleteness. Leaves of the reference tree *R* that are absent from the query tree *T* do not create a problem as they can be easily pruned from *R* without affecting the optimization problem. Leaves in *T* missing from *R* need care. Appendix A.4 details how missing data are handled in our approach, and Figure S2 provides evidence of its effectiveness.

### Application 1: Outlier in gene trees using MCOA

The phylogenomic analysis pipeline introduces various forms of error into gene trees, compounding biological processes such as ILS that create real gene tree discordance. Some errors leave subtle traces, while others distort the inferred trees more dramatically by introducing abnormally long branches or drastically altering the tree topology. These errors can impact downstream analyses (Laurin-Lemay et al., 2012; Philippe et al., 2017; Salichos and Rokas, 2013; Springer and Gatesy, 2018). As a result, several efforts have been made to detect anomalies in gene trees automatically. Mai and Mirarab (2018) proposed the TreeShrink method to identify species with unexpectedly long branches in a tree. Gori et al. (2016) proposed clustering gene trees using various distance metrics. Going further, de Vienne et al. (2012) introduced Phylo-MCOA to use multiple co-inertia analysis (MCOA) to embed all gene trees into a single Euclidean space such that pairwise distances of leaves match those of the input trees as much as possible. Using the principal coordinates of this space enables the detection of outlier genes and species in a two-step process employing Tukey’s method (Hubert and Vandervieren, 2008) followed by more complex steps. A recent implementation of Phylo-MCOA is available in an R package called PhylteR (Comte et al., 2023).

We propose to use TCMM to improve PhylteR outlier detection (Fig. 1a). We examine an example based on the simulated 100-taxon dataset (see *Datasets*) to motivate the approach (Fig. 1b-d). A species tree can have rate variation across branches, including highly diverged clades (Fig. 1b). Additionally, even in the absence of error, individual gene trees can have highly discordant topologies (e.g., Gene 17, Fig. 1b) or branch lengths (e.g., Gene 16) due to heterotachy (Lopez et al., 2002). These differences can make pairwise distances across different gene trees dissimilar and render embeddings noisy (Fig. 1c). In particular, branch length variations can lead to outliers even absent any topological discordance (e.g., Gene 16 in Fig. 1c). TCMM, by matching gene trees to a shared metric space, reduces the distance of the embeddings to the median (consensus) embedding (Fig. 1d). The more concentrated distributions have the potential to better reveal *topological* outliers. Moreover, *λ* defines the degree to which gene trees are matched to the shared space, allowing us to identify different types of outliers. With *λ* = 0, we match gene trees maximally to each other, reducing the noise created by rate heterogeneity both across and within gene trees, resulting in more concentrated embeddings and revealing outliers that would otherwise be obscured by noise in the data (e.g., Gene 17 becomes an outlier with *λ* = 0 in Fig. 1c). As *λ* increases, the ratios between original branch lengths are increasingly maintained (average rate heterogeneity across gene trees is reduced by TCMM). As a result, outliers with unexpectedly long branches are detected (e.g., Gene 16 in Fig. 1b,c). A *λ* value in between the two extremes trades off the two aspects.

**Figure 1.**
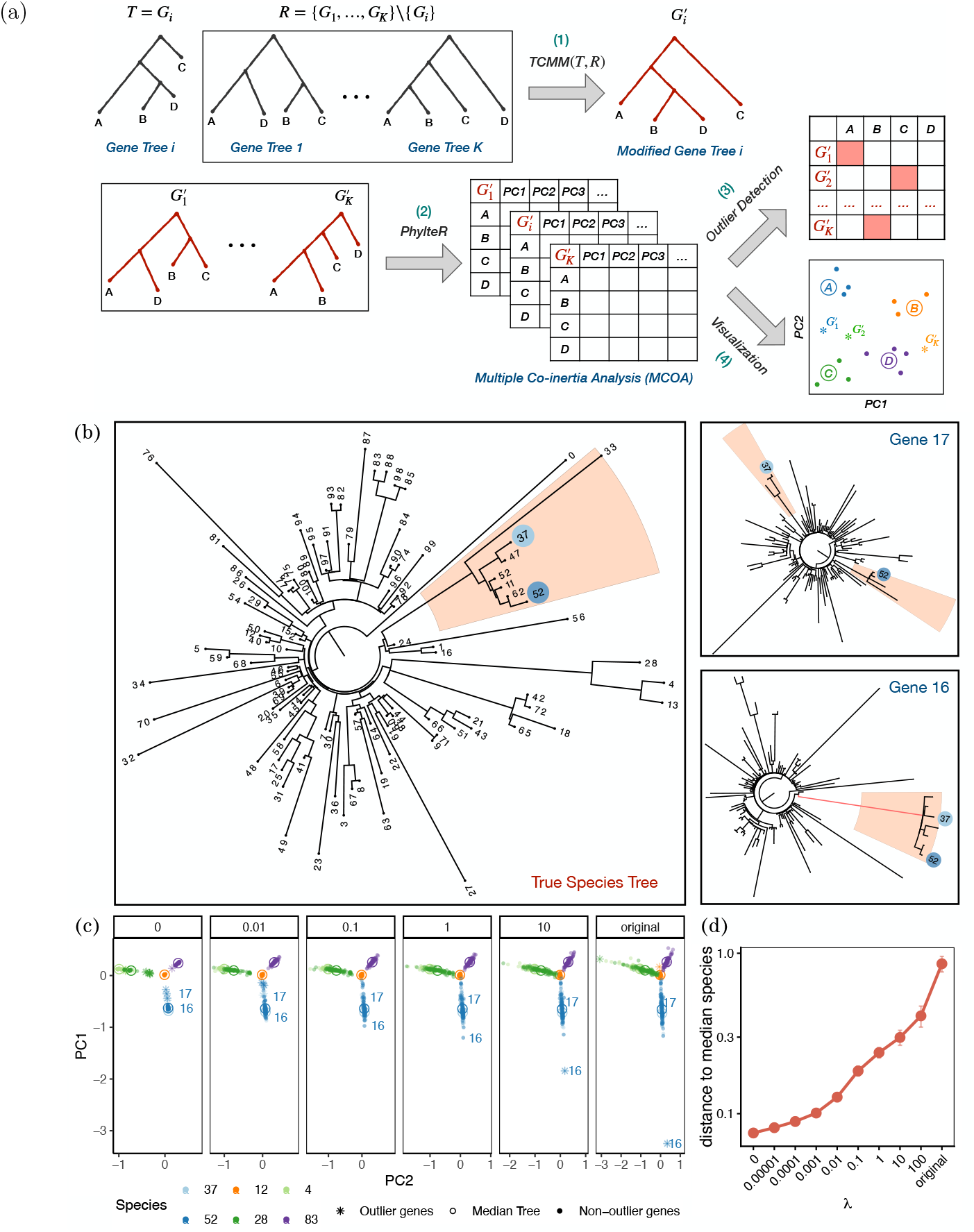
(S100-default Dataset) (a) The TCMM pipeline for outlier detection. Each modified gene tree 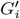 is the output of running TCMM with the original gene tree *G*_*i*_ as the query and the rest of the gene trees as the reference trees. The modified gene trees are then fed into the PhylteR pipeline, where the multiple co-inertia analysis (MCOA) is performed. Lastly, the principal coordinates (PCs) for each modified gene tree are used for the outlier detection and visualization tasks. (b) An example of the outliers detected by TCMM+PhylteR for one replicate. Left: the true species tree with the two species 37 and 52 and their corresponding subtree highlighted. Right: two outlier genes (16 and 17) for these species. Gene 17 is an outlier due to discordance from the true species tree topology, whereas gene 16 is an outlier due to a long branch (marked in red) separating species 37 and 52 from other species. (c) The PCA plot shows the projections of six species onto the first and second principal coordinates (PCs) with different regularization coefficient (*λ*) values. The position and the status (outlier or non-outlier) of genes 16 and 17 are marked for species 37 and 52 on each plot. (d) Comparing the Euclidean distance to the median species for true gene trees. (See also Fig. S3).

We evaluate TCMM’s performance on both simulated and biological datasets, where gene trees have branch lengths measured in units of expected substitutions per site (SU). We apply the unweighted consensus version of TCMM to each gene tree, using it as the query tree and treating the remaining gene trees as reference trees. The modified gene trees are then fed into the PhylteR pipeline to generate Euclidean embeddings, which are used for visualization and outlier detection. For simplicity, we skip the second step of outlier detection used in Phylo-MCOA and report only the outliers identified through Tukey’s method. We apply this procedure with and without regularization and examine how varying *λ* ∈ {0, 1e−5, 1e−4,…, 10, 100} influences the outliers detected. The same process is repeated with the original gene trees without running TCMM. To measure the effectiveness of outlier detection, we calculate the improvement in the normalized RF (nRF) distance from gene trees to the species tree *S*, compared to changes in nRF by random removal of the same number of taxa, leading to a metric called ΔRF (see Appendix A.2). We also use the normalized quartet score (nqs), defined as the proportion of shared quartets between two trees, to quantify the similarity between filtered gene trees and the species tree. We use the true species tree when available (simulated datasets) and the ASTRAL-III inferred species tree otherwise. We quantify whether outliers are the same as those found by the branch length-based method TreeShrink by reporting the containment Jaccard similarity (Appendix A.2) between the outliers detected by TCMM +PhylteR and those detected by TreeShrink. To evaluate the effect of outlier removal in downstream analyses, we use the filtered gene trees to infer the species tree using ASTRAL-IV (Zhang et al., 2025). For topology, we infer a species tree from the filtered gene trees and compare it to the true species tree using normalized RF distance (nRF). We also assess the accuracy of CU branch lengths, which are a direct function of gene tree discordance (Sayyari and Mirarab, 2016). In this analysis, we fix the species tree topology to the true topology and use the filtered gene trees to estimate CU branch lengths, expecting underestimation due to errors. We report mean absolute error (MAE) and bias, defined as the mean difference between estimated and true branch lengths.

### Application 2: Species tree branch lengths

Two-step methods of species tree inference produce a species tree topology from a set of gene trees. To be useful for downstream analyses (e.g., dating), the species tree requires branch lengths. Scalable and accurate methods such as ASTRAL (Mirarab et al., 2014b) and NJst/ASTRID (Liu and Yu, 2011; Vachaspati and Warnow, 2015) produce only topology. We can assign coalescent unit (CU) branch lengths for internal branches of the species tree using gene tree topologies alone, but this approach can lead to systematic underestimation (Forthman et al., 2022; Sayyari and Mirarab, 2016), and the lack of terminal branch lengths makes it less useful. Since gene trees have SU branch lengths and many downstream analyses require SU lengths, we can seek to transfer the SU branch lengths from gene trees onto the species tree. This should be possible because gene tree branch lengths are a function of species tree SU branch lengths (Fig. 2a). Briefly, we can assume each branch of the species tree has a substitution rate, and gene tree branches inherit these rates as well as per-locus rate multipliers (Mallo et al., 2016).

**Figure 2.**
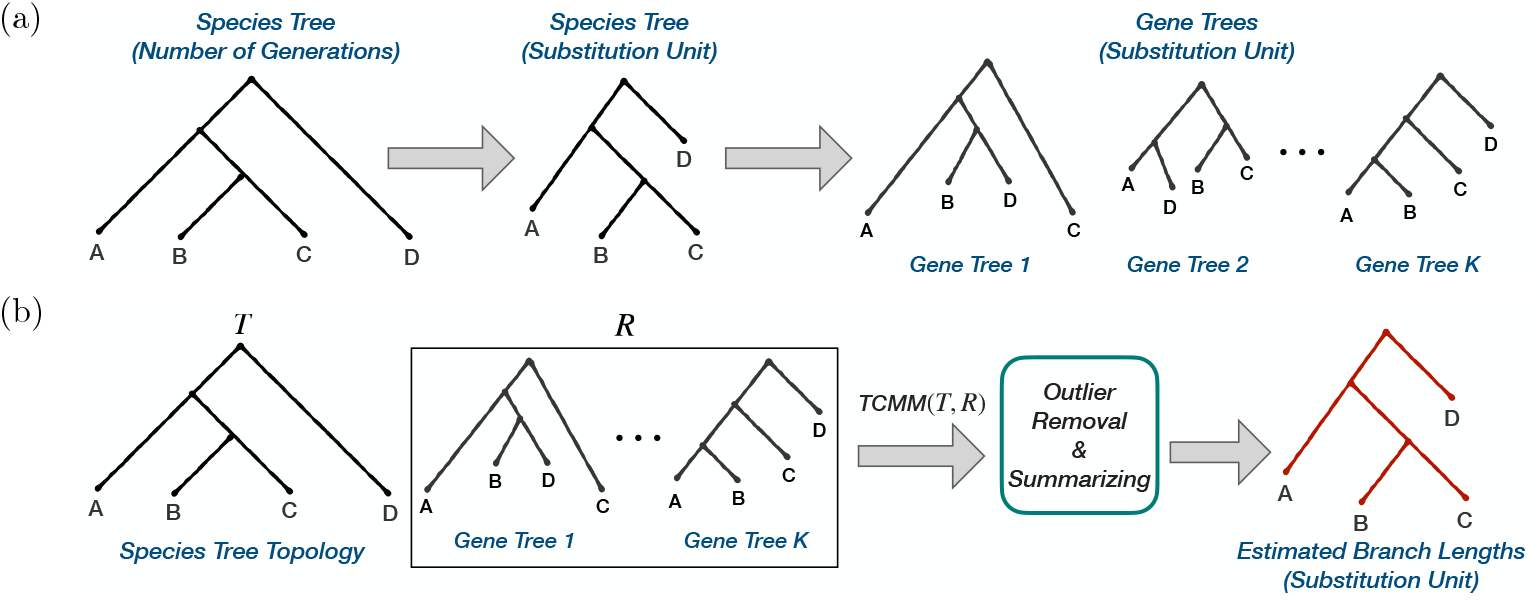
(a) Assumed data generation model, where species tree branch lengths measured in the unit of generations is multiplied by per-branch mutation rates to obtain SU length. Gene trees inherit these rates, with further per-locus variation. (b) The TCMM pipeline for branch length estimation with the species tree topology as the query and the gene trees with branch lengths in substitution unit (SU) as the reference trees. The final output is the species tree with SU branches.

Binet et al. (2016) proposed a method called ERaBLE that uses a least squares optimization applied to patristic distances of the gene trees to estimate species tree lengths, effectively taking a weighted average. ERaBLE allows for rate heterogeneity only across loci. It estimates a single scalar rate per gene tree by matching them simultaneously to the species tree. Tabatabaee et al. (2023) showed that ERaBLE overestimates terminal branches because it ignores deep coalescence. They propose a coalescent-based method called CASTLES to explicitly model the differences between the species and genic divergence times. While CASTLES and its newer version CASTLES-Pro are more accurate than ERaBLE in MSC simulations, their accuracy can decrease under high levels of HGT (Tabatabaee et al., 2025). This observation raises the question: Can a flexible model-free method assign SU lengths to the species tree while attaining high accuracy when gene tree heterogeneity is not solely due to ILS? Here, we use TCMM to summarize gene tree SU branch lengths onto the species tree (Fig. 2b). Unlike ERaBLE, which estimates only one scalar parameter per gene tree, we assign one parameter for each branch of each gene tree. Counterintuitively, we use gene trees as the reference and the species tree as the query in one of two ways. One approach, called consensus, uses Problem 3 to jointly match the species tree (as query *T*) to multiple gene trees (as the reference *R*). Here, we simply set all the *α*_*k*_ weights to 1. Our second (and default) approach is to run TCMM per gene (i.e., Problem 1), and summarize results at the end. Results provide us with a set of estimated branch lengths 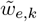 where *e* ∈ *E*_*S*_ and *G*_*k*_ ∈ *G*. These lengths can then be further summarized into a single value 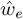 for each branch using common averaging methods — arithmetic mean, harmonic mean, and geometric mean. When using arithmetic means, this approach is similar to using the consensus method; Remark 2 (supplementary material) shows that the two are identical if all output lengths are strictly positive. Despite this (near) equivalence, the per-gene approach offers the benefit of enabling outlier removal. We remove both extremely small outliers introduced by TCMM and large outliers, which can arise due to gene tree errors, using the procedures detailed in Appendix A.3. We address missing data using the approaches noted in Appendix A.4.

We can pair TCMM with other branch length estimation tools. In this approach, we take the species tree with branch lengths estimated by another tool (e.g., CASTLES-Pro (Tabatabaee et al., 2025)), and apply the regularized version of TCMM (Problem 2) to this tree. This allows us to refine the input while preserving its branch lengths to an extent controlled by *λ*. One challenge is selecting the parameter *λ*. Here, we use an automatic approach to select *λ* from a predefined set of options. We do this by choosing the *λ* that minimizes the sum of the estimated species tree branch lengths. This choice is rooted in the minimum-evolution principle (Rzhetsky and Nei, 1993) used by a family of distance-based approaches.

We evaluate TCMM for this application using both simulated and real datasets. On simulated data, the model species tree branches have true SU lengths (*w*) obtained by multiplying their lengths (number of generations) with varying mutation rates, creating non-ultrametric trees. We compare the true *w* values to estimated values 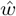 to quantify the mean absolute error 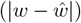 denoted MAE, absolute logarithmic error 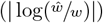, and bias 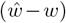, averaged across all branches of each species tree after excluding the outgroup. We compare TCMM to four leading methods studied by Tabatabaee et al. (2025), all run on the true species tree topology. We include CASTLES-Pro, which accounts for coalescence times predating species divergence times. We examine the commonly used method of running RAxML (Stamatakis, 2006) on concatenated gene alignments to estimate branch lengths of a fixed species tree (referred to as Concat+RAxML in our experiments). We also include two methods that like TCMM use patristic distances. A simple alternative is averaging patristic distance matrices across gene trees and using a distance-based method (FastME, Lefort et al., 2015) to assign branch lengths to *S* (called Patristic(AVG)+FastME in our experiments). A more advanced version of this idea is ERaBLE (default), mentioned earlier.

### Simulated Datasets

For application 1, we studied two simulated datasets, one with introduced errors and one without, and five biological datasets. For application 2, we studied four simulated datasets with different levels of gene tree estimation error, ILS, horizontal gene transfer (HGT), and rate heterogeneity. We also studied three microbial datasets. All biological datasets are detailed in the results section.

#### S200-perturbed dataset

We started from a 201-taxon Simphy-simulated dataset originally published by Mirarab and Warnow (2015) and perturbed its sequences to introduce errors to study Application 1. We focused on the condition with medium ILS (i.e., 2 million generations and a speciation rate of 1*e* − 7), which contains 50 replicates, each with 1000 gene trees. We introduced errors as follows. For each replicate, we perform *N* + 1 rounds of perturbation, with *N* ∼ Poisson(3), leading to at least one round and four rounds on average. In each round 0 ≤ *r* ≤ *N*, we select *k*_*r*_ ∼ Poisson(*λ* = 40) genes to be modified uniformly at random, and choose the proportion of taxa to be perturbed by drawing *t*_*r*_ ∼ Beta(10, 90) (mean: 0.1, sd: ≈ 0.03). Instead of selecting these perturbed (i.e., erroneous) species (ℰ _*r*_) uniformly at random, we use a random sampling strategy that encourages ℰ _*r*_ to be closer to each other on the tree than a uniformly random sampling would lead to. To do so, we root the species tree on a randomly chosen taxon and repeatedly traverse from the root to leaves, taking left/right children at random, and add any observed leaf to the set ℰ_*r*_; we repeat until |ℰ_*r*_| = *r*200 *× t*_*r*_*1*. To perturb these selected taxa, we first randomly reroot the species tree. Then, for each perturbed gene 1 ≤ *i* ≤ *k*_*r*_, we select a “source” taxon *s*_*r*,*i*_ using the same random sampling process used to select *ℰ* _*r*_, and rerooting the tree at this source between genes. For each gene *i*, we then draw *ρ*_*r*,*i*_ ∼ *Beta*(*α* = 16,*β* = 13) (mean 0.55, sd: 0.09) and replace the first *ρ*_*r*,*i*_ portion of each target sequence in ℰ_*r*_ with the same sites from *s*_*r*,*i*_ in gene *i*. Thus, each target sequence becomes a chimera between the *s*_*r*,*i*_ and its original sequence. This procedure leads to zero to 81 perturbations per gene (3.2 in expectation: 4 *×* 40 *×* 0.1 *×* 200 *×* ^1^*/*_1000_). We then re-infer a gene tree using FastTree-II (Price et al., 2010), given these perturbed sequences. We removed one replicate because the estimated gene trees (prior to introducing errors) contained numerous large polytomies. For each replicate, we used the first 100 gene trees in our analyses to reduce the accuracy of species trees and make the impacts on them more visible.

#### S100 simulated dataset

We reused the published 101-taxon 1000-gene dataset (S100-default) simulated by Zhang et al. (2018) using Simphy (Mallo et al., 2016), which Tabatabaee et al. (2023) then updated to include species trees with SU branch lengths (see Table S1 for all parameters). Our earlier results, shown as motivation for Application 1, use this dataset, and we use it in Application 2 as well. To recap the simulation procedure, briefly, ultrametric species trees with generation times are simulated using a birth-death model; to get SU length, each branch length is multiplied by a fixed mutation rate across the tree (drawn from a LogNormal distribution with mean 4 *×* 10^−8^) and a rate multiplier drawn from a Gamma distribution (mean: 1; inverse of variance, *α*, drawn from a LogNormal(1.5,1), leading to variance ranging in our replicates between 0.0153 to 2.6027). The 50 replicates of this dataset are highly heterogeneous in terms of ILS, with the mean nRF distance between the model species tree and the true gene trees at 46%. Gene trees were estimated using FastTree-2 (Price et al., 2010) from 200bp, 400bp, 800bp, and 1600bp alignments, leading to average nRF distance between true and estimated gene trees of 55%, 42%, 31%, and 23%, respectively. In application 1, we only use the first 100 gene trees for each replicate due to computational constraints and to make differences between methods more pronounced.

To tease out the impact of ILS levels and rate heterogeneity modes in Application 2, we used the updated Simphy code to simulate six model conditions of the same S100 dataset (naming them S100-truegt), each condition with 50 replicates (Table S1). In these analyses, intended for validation and sanity checks, rather than biological realism, we used true gene trees; therefore, no sequences were generated. Simphy parameters were kept the same, except for the following changes. For varying ILS, we created three model conditions by changing the population size (low, med, high ILS: *N*_*e*_ = 4e+4, 4e+5, and 4e+6, resp.) and turned off lineage-specific rate variation so that the species tree becomes ultra-metric in SU units. Note that each condition still retains some level of difference in ILS levels across replicates due to differing tree heights. The average nRF distance between true gene trees and the true species tree for low, med, and high ILS model conditions is 8%, 44%, and 89%, respectively. For studying the impact rate heterogeneity, we created three conditions. We refer to the original simulation as `hl+hg+hs`; this condition has rate multipliers per species, per gene, and per individual branches of individual gene trees. We added `hl+hg` which has a rate multiplier per gene and one multiplier per each branch of the gene tree (`hl+hg`); however, there is no global per species rate, making the species three ultrametric. We also added `hl+hs`, which assigns a rate multiplier per species tree branch, making the tree non-ultrametric, and also assigns a rate per gene.

## Results

### Application 1: topology focused MCOA

#### S200-perturbed Simulated Dataset

Using TCMM with PhylteR increases the accuracy of error detection compared to using PhylteR alone for some values of *λ* (e.g., 0.1 and 1), as reflected in higher true positive rates (TPR) and improved ROC curves (Fig. 3a). The PhylteR threshold *k* allows us to control sensitivity, with lower *k* corresponding to increased total number of outliers. Across all *k*, the Area Under the Curve (AUC) increases from 0.811 with PhylteR alone to 0.884 with PhylteR+TCMM with *λ* = 0.1. However, while patterns of performance remain similar across *k*, choices of *k* with low false positive rate (FPR) are of particular interest. As default, we use *k* = 1.5, which provides a low FPR and substantial TPR (e.g., FPR = 0.008 and TPR = 0.425 for *λ* = 0.1). On this dataset, TreeShrink (with default threshold) has a substantially lower TPR and higher FPR (FPR = 0.026 and TPR = 0.006).

**Figure 3.**
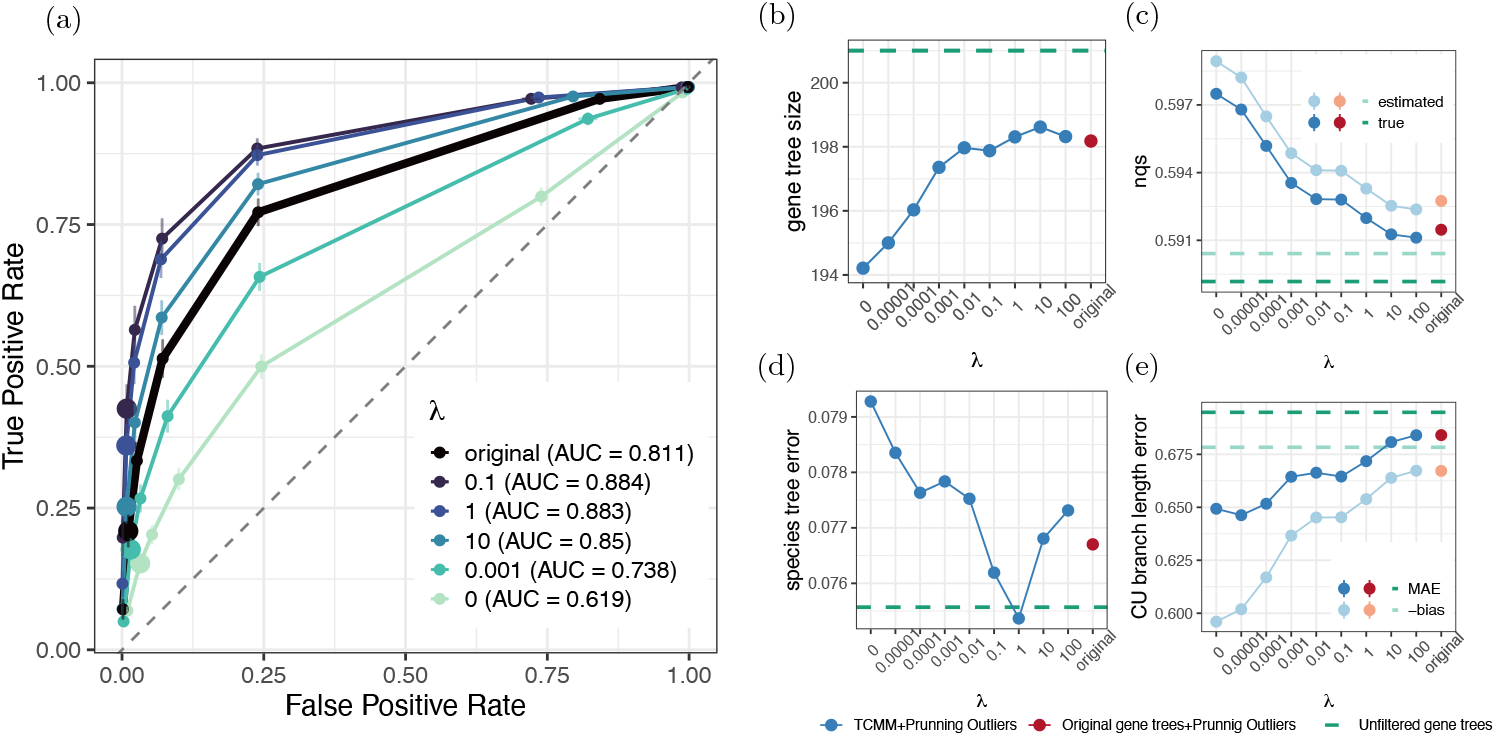
(S200-perturbed Dataset) (a) ROC curve for PhylteR+TCMM outlier detection as we change its threshold *k* (dots). Colors show different inputs to PhylteR. Black shows the original gene trees, whereas other colors are TCMM gene trees with different *λ* values. *k* = 1.5 is marked with a larger size. Dashed line shows random outlier removal. Area under the curve (AUC) values are shown in the caption. (b) Number of remaining taxa in gene trees after outlier removal. (c) Normalized quartet score of pruned and unfiltered gene trees to the true and estimated species trees. Above the green line shows improvement compared to unfiltered gene trees. (d,e) Species tree accuracy after outlier removal, showing nRF distance of estimated tree topology (d) and mean absolute error (MAE) and bias of estimated coalescent unit (CU) branch lengths (e) to the true species tree. Below the green line shows improvement compared to unfiltered gene trees. Similarity of MAE and bias indicates that the error is due to a systematic under-estimation of lengths, which is reduced after filtering.

Comparison to PhylteR depends strongly on the choice of *λ*, with low values (e.g., *λ* = 0) showing worse performance than PhylteR alone, and very high values (e.g., *λ* = 10) showing only modest improvements over PhylteR. However, for the middle range (*λ* = 0.1 or *λ* = 1), the improvements are substantial. In this range, the number of removed outliers (roughly 3 per gene tree) is about the same as PhylteR without TCMM (Fig. 3b), showing that the improvements of TCMM are not due to finding more outliers but rather finding more erroneous sequences. With lower *λ* values, PhylteR+TCMM removes many more topological outliers (e.g., 6.8 with *λ* = 0), which leads to increased similarity between the outlier-pruned gene trees and both the true and estimated species trees (Fig. 3c). However, these reductions in discordance come at the cost of removing many more data points (Fig. 3b) and higher FPR (Fig. 3a). To test if the reduction in signal due to pruning by each PhylteR+TCMM variant is justified by their reduction in noise, we used the resulting pruned gene trees to estimate the species tree topology (Fig. 3d) and CU branch lengths (Fig. 3e). For topology, in most cases, removing outliers *increases* the species tree error in terms of nRF distance (Fig. 3d). The only case with (slight) improvement in topological accuracy is *λ* = 1, which is among the values that best trades off precision and recall. All the other settings, including PhylteR without TCMM, result in increased topological error in ASTRAL trees, perhaps because the ASTRAL topology has some level of robustness to errors, and the loss of signal due to mistaken removal of correct sequences (false positives) can outweigh the inclusion of some noise.

Unlike topology, CU branch lengths are highly sensitive to error (Sayyari and Mirarab, 2016) and thus improve consistently as we remove more outliers (Fig. 3e). Any level of filtering, with or without TCMM, reduced error, but the largest reduction was obtained by TCMM+PhylteR with the smallest *λ* values. The distinct patterns observed between topology and branch length reflect the higher sensitivity of CU branch lengths to gene tree error.

### Simulated Dataset (S100-default Dataset)

On the S100-default dataset, which contains no sequence errors (only the typical gene tree estimation error), both PhylteR and TCMM+PhylteR still detect some outliers. However, the number of outliers detected by TCMM+PhylteR on this dataset is substantially lower than on the S200-perturbed dataset (Fig. S3b). For example, with *λ* = 0.1, 0.11% of taxa are removed across all conditions of S100-default compared to 1.46% for S200-perturbed with the same *λ*. Pairing TCMM with PhylteR results in fewer removals for *λ >* 0.01 but not for lower *λ* (Fig. S3b). While removing these outliers tends to decrease discordance between the gene trees and the true species tree, these improvements are not substantial (no more than 2%), likely due to the small number of outliers (Fig. S3c). We observe that decreasing *λ* makes the outliers detected by TCMM+PhylteR increasingly dissimilar to those detected by TreeShrink, and the reduced emphasis on branch lengths leads to the detection of more topologically aberrant outliers (Fig. S4a), a pattern we referred to earlier with one example (Fig. 1b). Moreover, compared to randomly removing the same number of leaves, outliers detected by TCMM+PhylteR yield a greater reduction in nRF distance for lower values of *λ*, with this pattern observed across all levels of gene tree error and more pronounced in higher-quality gene trees. These observations are consistent with our expectation regarding the role of *λ* in shaping the type of outliers detected by our method (see *Methods*).

#### Biological Dataset

We next studied six biological datasets (mammals, frogs, plants, and two datasets on Xenacoelomorpha, Insects) adopted from Mai and Mirarab (2018). These datasets cover a range of number of species, number of genes, and evolutionary depth (Appendix A.1). Some patterns observed in the simulated datasets persist. For example, As *λ* decreases, MCOA embeddings become more concentrated and closer to the median embedding for each species (Fig. S5). Other patterns differ from the simulated datasets. The impact of *λ* on the number of outliers depends on the dataset, with some, like mammals, behaving like simulated data and others, like insects and plants, showing a reduction in the number of outliers after applying TCMM (Fig. 4a). The total number of outliers also varies across datasets, ranging from 0.01 on the Xenacoelomorpha dataset of Rouse et al. (2016) (*λ* = 1) to 3.16 for the Frogs dataset (*λ* = 0.001).

**Figure 4.**
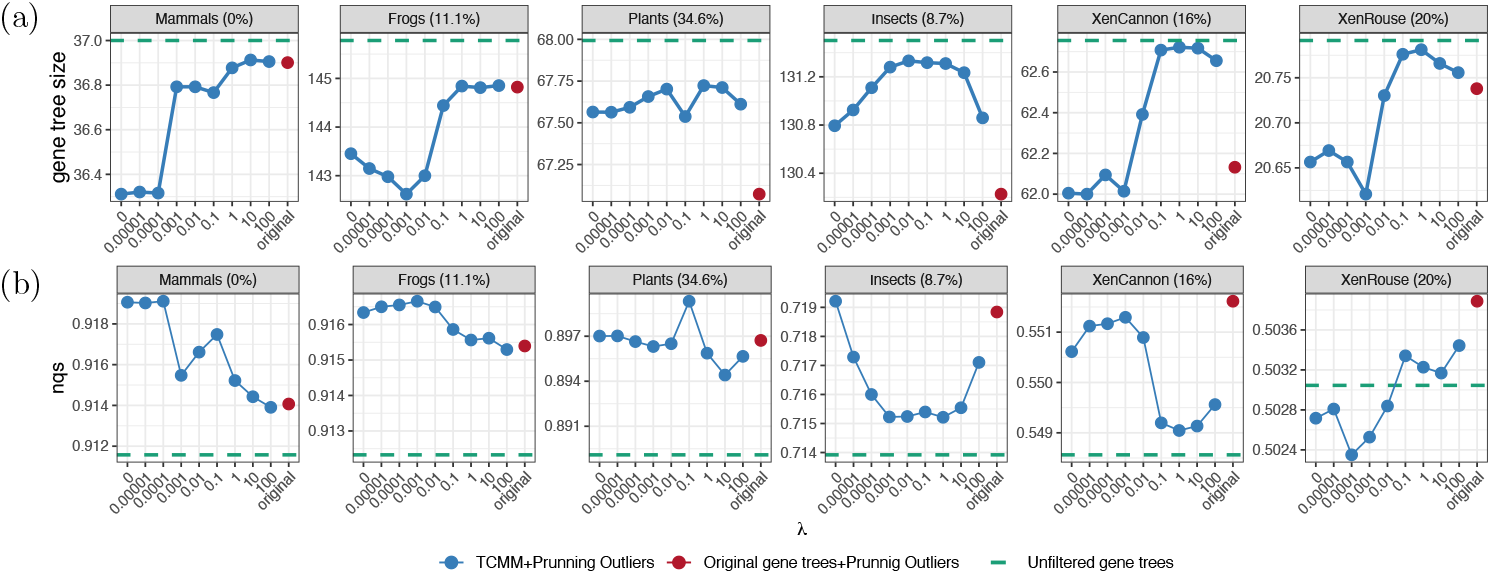
(Biological Dataset) (a) Number of remaining taxa in gene trees after outlier removal for each *λ*. The amount of missing taxa from each dataset is written as a percentage. (b) Normalized quartet score to the ASTRAL estimated species tree. The unfiltered original gene trees are shown in green.

Removing outliers identified by TCMM+PhylteR generally increases the similarity to the estimated species tree for most, but not all, datasets (Fig. 4b). The amount of improvement tends to be higher when the original gene trees are less discordant with the estimated species tree. In particular, for mammals and frogs, which had low discordance, TCMM+PhylteR improves quartet scores more than using PhylteR alone with any *λ*, and more so with lower *λ* values. For plants and insects, some *λ* values are better than PhylteR alone, while others are not, but all cases lead to improved quartet scores compared to no filtering. In the case of the Xenacoelomorpha datasets of Cannon et al. (2016), pairing TCMM with PhylteR can reduce improvements compared to PhylteR alone though it still results in improvements. The Xenacoelomorpha dataset of Rouse et al. (2016), however, was different; removing outliers makes the gene trees more dissimilar to the species tree with TCMM+PhylteR but not with PhylteR alone or high *λ*. The gene trees in this dataset have the highest discordance with the estimated species tree among all the studied biological datasets (Fig. 4b), suggesting that gene tree quality and species tree accuracy may impact the effectiveness of outlier detection using TCMM.

## Application 2: Species tree branch lengths

### Simulated Data

#### Choosing a version of TCMM

The per-gene approach with arithmetic mean exhibits lower bias compared to the consensus version of TCMM without compromising on MAE (Fig. S6a), making it the preferred choice. Another advantage of the per-gene approach is its ability to perform outlier detection and removal, effectively eliminating extremely small estimated branch lengths that are often present in the consensus approach. In fact, the consensus method can assign branch lengths that are essentially zero, particularly for smaller *λ* values. However, these underestimations are mitigated in the per-gene method due to outlier removal (Fig. S6bd). Incorporating outlier removal also reduces bias and MAE in terminal branches while slightly increasing MAE for internal branches (Fig. S6a). The consensus approach of TCMM is the most similar to the distance-based method ERaBLE, both in methodology and performance (Fig. S6c). Thus, the increased performance compared to ERaBLE is primarily attributed to the combination of per-gene branch length estimates and outlier detection.

While all three methods (consensus, per-gene, and per-gene with outlier removal) perform similarly when initial branch lengths are available, the per-gene method with outlier removal is adopted as the default version of TCMM due to its performance in the absence of initial branch lengths and the advantages of outlier removal in biological datasets. Furthermore, examining varying *λ* reveals that as *λ* values increase, branch lengths estimated by the default TCMM method become progressively more similar to the initial branch lengths (CASTLES-Pro) in terms of their relative values (Fig. S6bd).

#### Comparison to other methods

On the S100-truegt dataset, when ILS levels are kept low, all methods run on the true gene trees have near-perfect accuracy (Fig. S7). However, when ILS increases, the error goes up for all methods, leading to TCMM outperforming other distance-based methods but not CASTLES-Pro. Focusing on medium ILS levels and changing modes of rate heterogeneity does not change the relative accuracy of methods (Fig. S8). When comparing conditions with ultrametric species trees (`hl+hg`) to non-ultrametric cases (`hl+hs`), we observe increased error for all methods, but not to the same degree. TCMM ‘s normalized MAE increases by 10%, while ERaBLE and AVG+FastME show increase by 27% or more (comparisons to `hl+hs+hg` are similar). Thus, TCMM is more robust to lineage-specific rate heterogeneity than other distance-based methods, likely because methods such as ERaBLE account primarily for genome-wide rate variation.

On the S100-default dataset (with heterogeneous ILS, simulated sequences, and estimated gene trees), TCMM is better than both the commonly used concatenation method (Concat+RAxML) and distance-based methods (ERaBLE and AVG+FastME) but not the coalescent-based CASTLES-Pro in terms of overall mean absolute error and bias (Fig. S9). The improvements are due to much better terminal branch calculation; for internal branches, TCMM has a slight disadvantage compared to both distance-based methods (ERaBLE and AVG+FastME), but not concatenation. Combining CASTLES-Pro with TCMM (using automatic *λ*) is substantially better than TCMM alone, yet it does not outperform CASTLES-Pro. This result confirms that CASTLES-Pro, which is specifically designed and optimized for ILS, outperforms cause-agnostic distance-based methods when gene tree discordance is dominated by ILS.

The patterns shift in simulations with HGT, where the best method is combining TCMM and CASTLES-Pro with automatic *λ* selection (Fig. 5). The estimation error of all methods increases as the rate of HGT increases, especially for the two highest HGT rates. CASTLES-Pro remains better than TCMM alone and other methods. TCMM is better than methods other than CASTLES-pro for terminal branches and slightly worse for internal ones. All methods are biased, and these biases increase as the rate of HGT increases. CASTLES-Pro has an underestimation bias, while the other methods have an overestimation bias, which is concentrated on terminal branches. TCMM has the lowest overestimation bias among methods other than TCMM+CASTLES-Pro (e.g., on average 50.4% lower bias than concatenation).

**Figure 5.**
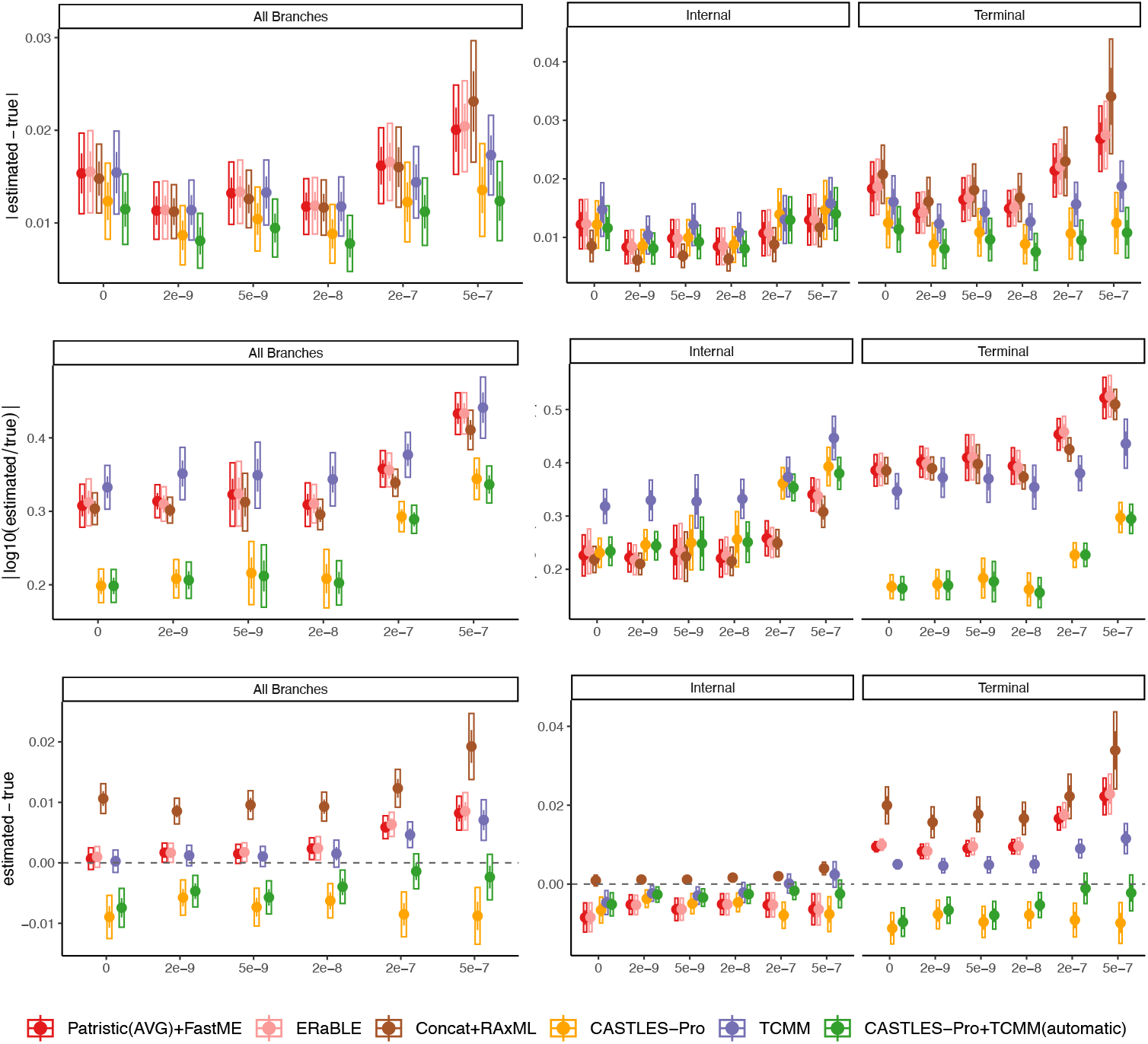
(HGT Dataset) Comparing TCMM with the alternative methods across various rates of HGT (x-axis). TCMM is run with *λ* = 0 (TCMM) and automatic *λ* selection (CASTLES-Pro+TCMM). The methods are compared using two error metrics (mean absolute error and mean absolute log error) as well as bias.

On this dataset, however, TCMM+CASTLES-Pro has the best accuracy for all measures of error and bias. On average, the mean absolute error of CASTLES-Pro+TCMM is 8.2% lower than CASTLES-Pro and 27.3% lower than TCMM alone. The clearest advantage is the reduction in bias, particularly at higher HGT rates: While TCMM reduces bias by 26.7% compared to CASTLES-Pro across all HGT rates, the TCMM+CASTLES-Pro combination achieves reductions of 27.3% and 42.1% over TCMM alone at HGT rates of 2 *×* 10^−7^ and 5 *×* 10^−7^, respectively. These results establish TCMM+CASTLES-Pro as a reliable choice for running TCMM across all tested datasets and model conditions (Fig. S10).

No single *λ* consistently outperforms others across all settings (Fig. S10). This variability makes it useful to select a *λ* automatically, eliminating the need for manual tuning. Automatic *λ* selection tends to choose larger *λ* values (Fig. S11), likely due to TCMM’s overestimation bias, which causes smaller *λ* values to result in overall higher estimated branch lengths. Note that even with high *λ*, CASTLES-Pro+TCMM benefits from outlier detection and removal, a feature absent in CASTLES-Pro alone.

### Biological data: AB branch length

We analyzed several microbial datasets, focusing on the separation between bacteria and archaea (AB branch). Moody et al. (2022) claimed that lower divergences obtained by some recent studies (Petitjean et al., 2015; Zhu et al., 2019) were the result of highly abundant HGT in microbial datasets (Arnold et al., 2022). They compared results with two alternative datasets of core genes, with less propensity for HGT. Following them, we reanalyzed a 72-taxon tree (Williams et al., 2020) with 49 core genes, including ribosomal proteins and other conserved genes, and a 108-taxon dataset (Petitjean et al., 2015) with 38 genes that only include non-ribosomal proteins. We compared results with a reanalyses of the microbial dataset by Zhu et al. (2019) with 10,575 species sampled evenly from bacteria and archaea and 381 genes (including many non-core genes). The species tree used in this analysis was inferred using ASTRAL and is rooted on the branch separating bacteria from archaea. Gene trees in this dataset lack 28% of taxa on average. On all datasets, we compare the length of the AB branch inferred using TCMM, CASTLES-Pro, and concatenation, all drawn on species topologies built using ASTRAL-III (Zhang et al., 2018) from the two gene sets. On these datasets, ERaBLE fails to return an output, and AVG+FastME produces an estimated tree with several long negative branches around the root; thus, we exclude these methods.

Using 49 core genes (including ribosomal genes), CASTLES-Pro, TCMM, and CASTLES-Pro+TCMM (automatic *λ* selected as 1) estimate a length of 1.79, 1.94, and 1.91 SU for the AB branch, respectively, compared to 1.43 SU for concatenation. In contrast, on the 38 non-ribosomal genes (presumably with more HGT), the AB branch is much shorter, as noted by Moody et al. (2022). TCMM and CASTLES-Pro+TCMM (with *λ* selected as 100) estimate AB branch length to be 0.87 and 0.76, respectively, which are higher than concatenation (0.43) but lower than CASTLES-Pro (1.05). The observation that both gene tree-based methods yield higher branch lengths than concatenation for non-ribosomal genes is consistent with Moody et al. (2022)’s findings that concatenation can underestimate branch length in the presence of HGT. The substantial difference between ribosomal and non-ribosomal genes observed using methods that account for gene tree discordance confirms the fundamentally different patterns of evolution between the two. In per-gene mode, TCMM estimates a separate branch length for each gene tree (Fig. 6 a,b). An examination of the distribution reveals substantial diversity among genes, with some indicating an AB branch length of more than 4 SU, while others suggest a length below 0.5 SU.

**Figure 6.**
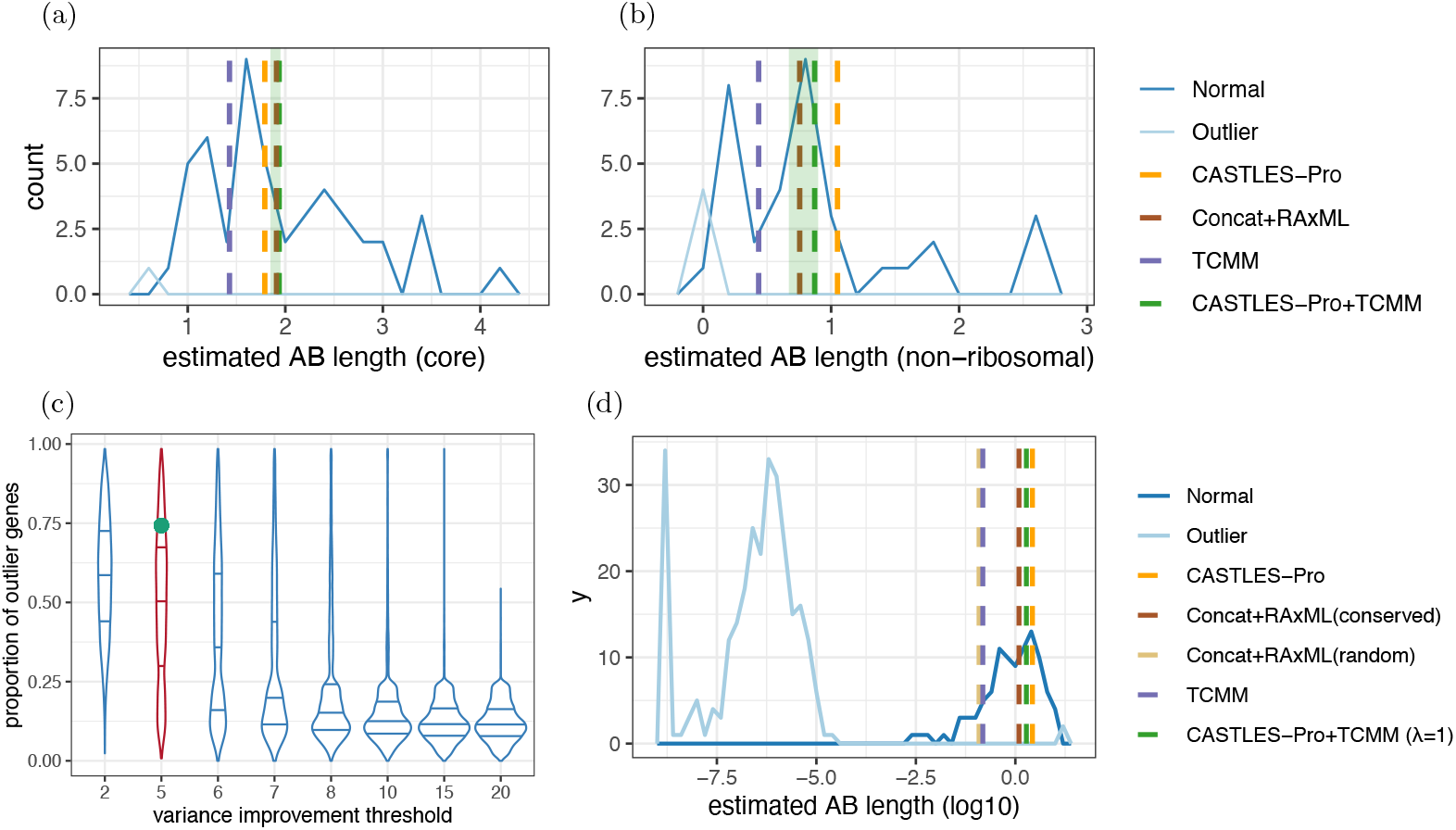
(a), (b) The distribution of the TCMM (*λ* = 0) estimated length for the branch separating bacteria and archaea per gene tree. Vertical dashed lines show the estimates of TCMM, CASTLES-Pro+TCMM, CASTLES-Pro, and concatenation. The range of the estimated branch lengths for the AB branch by CASTLES-Pro+TCMM for different values of *λ* is shown in green. (a) Uses the 49 core genes, with *λ* = 1 automatically selected for CASTLES-Pro+TCMM; (b) uses the 38 non-ribosomal genes, with *λ* = 100. (c), (d) (Zhu et al. (2019) Dataset). (c) The distribution of the outlier genes removed by TCMM (*λ* = 0) per branch for different variance improvement thresholds; the default threshold (5) and the AB branch under this threshold are highlighted in red and green, respectively. (d) The distribution of the TCMM estimated AB branch length per gene tree.

On the microbial dataset by Zhu et al. (2019), CASTLES-Pro, TCMM, and CASTLES-Pro+TCMM (*λ* fixed to 1 due to high running time) estimate branch lengths of 2.68, 1.89, and 1.24 SU, respectively, for the AB branch. As a result, both CASTLES-Pro and TCMM estimate drastically longer branch lengths than concatenation, which estimates 0.15 SU when selecting 100 random sites per gene and 0.12 SU when selecting the 100 most conserved sites (original paper selected 100 sites per locus due to computational limitations of concatenation with *>* 10000 tips) Once again, the distribution of estimated AB branch lengths across individual gene trees shows significant variation, ranging from as low as 10^−9^ to over 18 SU for some gene trees (Fig. 6d). In computing the final estimate, the outlier removal step of the per-gene method removes 283 out of 381 estimates before summarization (281 low outliers and 2 high outliers) This high level of outlier detection may seem excessive, a topic we will return to in the discussions. On this dataset many branches have large numbers of outliers removed, presumably due to rampant HGT. Compared to all branches in the tree, the AB branch is among those with the highest proportions of outliers (Fig. 6c).

## Discussion

We introduce TCMM, a versatile framework for reconciling the branch lengths of a query tree to match one or more reference trees. At its core, TCMM provides a way to match tree metric spaces, thereby decreasing branch length discordance between trees without changing topological discordance. The underlying optimization problem builds on the least-squares formulation long used in distance-based phylogenetics, with added regularization terms. The regularization provides a way to adjust how far the adjustment goes. However, with or without regularization, the output branch lengths do not automatically have a biological meaning; instead, they are intended to serve as input in downstream applications. Our paper focuses on two applications: outlier detection and species tree branch length estimation. In both applications, combining TCMM with existing methods improves accuracy in many but not all conditions.

In the outlier detection application, we use TCMM to match gene trees to each other by harmonizing their metric spaces so that the remaining outliers reflect topological differences. Without such harmonization, embeddings of tree leaves might not align due to divergent rates of evolution in entire genes or some branches of certain genes, rather than errors in data. Our results show that pairing TCMM with PhylteR improves the accuracy of outlier detection compared to both PhylteR and TreeShrink, and that removing these outliers can improve downstream analyses – particularly in settings that are more sensitive to outliers, such as branch length estimation. The regularization term *λ* helps us control the extent to which cases of substantial rate variation are marked as outliers. For example, with true gene trees, where the only sources of discordance are ILS and rate heterogeneity, we saw that changing *λ* directly controls whether topological or branch length outliers are detected (Fig. S4a). When simulated sequence data had errors, and in biological data, anomalies may simultaneously affect both topology and branch lengths, blurring the distinction between the two (Fig. S4c). Thus, in practice, it may be helpful to examine a range of *λ* values. Beyond detecting errors in the data, this type of harmonization may also prove useful in other applications, such as detecting changes in rates of evolution. Note that TCMM outputs a multiplier for every branch of every gene tree, and examining the distributions of these multipliers may prove useful in detecting patterns of rate variation across the tree and across loci.

Our second application matches gene trees to a species tree instead of matching them together to transfer SU branch length from gene trees to the species tree. While TCMM is not as accurate as CASTLES-Pro (which is designed based on the MSC model) under ILS-only conditions, TCMM provides several benefits. Foremost, in its per-gene mode, TCMM provides a length *distribution* for each species tree branch. By outputting a multiplier per branch per gene, it allows for across-genes heterotachy (Lopez et al., 2002), a feature missing from other methods. ERaBLE uses only one mutation rate per gene, making it far less versatile. CASTLES-Pro models rate heterogeneity across the species tree and overall rate changes across genes; however, it does not estimate a per-gene rate for each species tree branch. Our biological results show evidence (Fig. 6) of heterotachy. Even when outliers are excluded, the remaining loci still provide a range of lengths for the AB branch and other branches across the microbial tree of life, pointing to heterotachy.

The second advantage of TCMM for species tree branch length estimation lies in outlier detection. With input gene trees generated using models assumed by CASTLES-Pro, we observe no advantage in pairing it with TCMM. In contrast, in the presence of HGT, which is not modeled by CASTLES-Pro, we observe a lower bias when CASTLES-Pro is followed by TCMM. HGT creates branch length patterns that can widely vary from those of HGT-free genes, a notion that has been used by some previous methods of HGT detection (e.g., Dessimoz et al., 2008). The outlier detection step can eliminate gene trees with widely incongruent topologies, which, when matched to the species tree, would otherwise lead to extremely short branch lengths. We observe many such cases (283 out of 381) for the AB branch length in the microbial dataset. By omitting these genes, TCMM+CASTLES-Pro is able to estimate a much higher AB length compared to concatenation. Labeling the majority of genes as outliers, as TCMM does for the AB branch, may seem counterintuitive: How can more data be classified as outliers than not? This point gets at the heart of the debate about the AB branch. By removing enough horizontally transferred genes, we can estimate high values for the AB length, as Moody et al. (2022) argued. Concatenation (or even TCMM without outlier removal) estimates much shorter lengths given a large number of marker genes. Is removing many (or most) genes biologically justifiable? The answer depends on how we interpret the branch lengths and what we use them for. In one view, despite HGT, there is an underlying tree (vertically inherited), providing the backbone of the network representing the full evolutionary history. Then, to find the number of substitutions on a branch of the tree, we should remove horizontally transferred genes, which add not just noise but an underestimation bias. Thus, we do not consider “outliers” merely as data points that deviate from the majority for unknown reasons. Rather, in this case, the high topological discordance due to HGT results in short estimated lengths on the backbone species tree (near zero for TCMM). Thus, the primary role of outlier detection (responsible for removing 281 out of 283 outliers for the AB branch) is to eliminate these near-zero estimates, which are prevalent due to the high levels of HGT around the branch. In a different view, if horizontal evolution is pervasive and concentrated enough to render the backbone tree an inadequate model, one might instead aim to estimate an overall genome-wide level of divergence between two groups of taxa (e.g., archaea and bacteria). In such cases, removing large parts of the genome is less defensible. Regardless, by providing a distribution per branch, TCMM offers a way to better represent the full evolutionary history. A promising direction of future research is to investigate whether these distributions can be incorporated into downstream applications such as UniFrac calculation (Lozupone and Knight, 2005).

A practical question arises: Should CASTLES-Pro be followed by TCMM for branch length estimation? The answer, based on our findings, is nuanced. If we believe the cause of discordance is mostly incomplete lineage sorting, the answer seems to be no. Additionally, TCMM in its current form is not equipped to handle multi-copy gene trees Chaudhary et al. (2013)— a potential avenue for future development. However, in scenarios involving single-copy gene trees with substantial levels of HGT or other aberrant sources of discordance (e.g., caused by hidden paralogy, alignment errors, or missing data), TCMM can enhance accuracy by detecting and removing outliers. Since homology errors abound in phylogenomic datasets (Springer and Gatesy, 2018; Steenwyk et al., 2023) and outlier loci can leave an outsized impact (Shen et al., 2017), running TCMM with CASTLES-Pro as input to detect the outliers seems prudent in most analyses.

The TCMM framework can have wide-ranging applications beyond those explored in this study. Once the species tree branch lengths are estimated and perhaps calibrated to the unit of time, TCMM can also be used to match gene trees to the species tree (opposite of our approach here) for downstream applications such as visualization (e.g., using Bouckaert (2010)), estimation of average rates, and perhaps even dating individual gene trees, topics that can be explored in the future. The linear algebraic approach employed here effectively compares two trees by matching one to the other. Unlike many commonly used phylogenetic metrics, this matching incorporates both tree topology and branch lengths. Thus, the results of the matching could potentially be translated into new metrics of tree distance and new median tree problems. Future research should investigate these metrics and whether they provide additional insights beyond those offered by existing methods.

## Acknowledgment

This work was supported by the National Institutes of Health (1R35GM142725) grant to S.M. This work used Expanse at San Diego Supercomputing Center through allocation ASC150046 from the Advanced Cyberinfrastructure Coordination Ecosystem: Services & Support (ACCESS) program, which is supported by U.S. National Science Foundation grants #2138259, #2138286, #2138307, #2137603, and #2138296.

## Code and Data availability

The software is available at https://github.com/shayesteh99/TCMM.git. Data is available on Github https://github.com/shayesteh99/TCMM-Data.git.

