## Supplemental material for "Branch Length Transforms using Optimal Tree Metric Matching"

**B Supplementary Tables** **7**

**C Supplementary figures** **9**

### A Supplementary text

#### A.1 Datasets

##### A.1.1 Biological datasets for application 1

**Mammals:** We studied a mammalian dataset by Song et al. (2012). This dataset contains 37 species (36 mammals and Chicken as outgroup) and 424 genes, where complete gene trees were inferred using RAxML in the re-analysis of the dataset by Mirarab et al. (2014a). Previous analyses (Gatesy and Springer, 2014; Mirarab et al., 2016) have identified several sources of error in this data, making it a good candidate for error detection.

**Xen-Cannon and Xen-Rouse:** We examined two datasets aimed at determining the phylogenetic position of Xenacoelomorpha, a phylum of bilaterally symmetrical marine worms, in the tree of life. The first dataset, referred to as XenCannon in our results, was studied by Cannon et al. (2016) and consists of 78 species and 213 genes. The second dataset, XenRouse, was studied by Rouse et al. (2016) and includes 26 species and 393 genes. The question of whether Xenacoelomorpha is the sister clade to the remaining Bilateria has been a long-standing debate (e.g., Philippe et al. (2011)), making these two datasets particularly significant for analysis.

**Frogs:** We analyzed a dataset published by Feng et al. (2017) that explores the evolutionary history and rate of evolution in frogs. This dataset includes 164 species, with 156 frog species and 8 outgroups, and is composed of 95 genes. For our analysis, we use the RAxML gene trees from this study.

**Plants:** We analyzed a plants and algae dataset published by Wickett et al. (2014), consisting of 104 species (including 4 outgroups) and 852 genes. Among the datasets used in our analysis, this one has one of the highest levels of missing taxa, with an average of 34.6% missing taxa.

**Insects:** We studied a phylotranscriptomic dataset of insects first published by Misof et al. (2014) to resolve the conflict around the evolutionary relationships between major insect orders. This datasets includes 144 species and 1478 genes. In our analysis, we use the RAxML estimated gene trees published by Sayyari et al. (2017).

#### A.2 Measurement metrics

##### A.2.1 Definition of ΔRF

Let $S$ be a reference species tree. On simulated datasets, we use the true species tree as $S$, while on biological datasets, we use the species tree inferred by ASTRAL-III. For a gene tree $G$, let $\mathcal{O}$ denote the outliers identified by a method. Let the $G \setminus \mathcal{O}$ denote a pruned version of tree $G$ removing leaves of $\mathcal{O}$ and suppressing degree 1 nodes. We randomly remove the same number of species removed by a method from the gene tree ($|\mathcal{O}|$) in 10 replicates ($\mathcal{R}_1 \ldots \mathcal{R}_{10}$, $|\mathcal{R}_j| = |\mathcal{O}|$). The improvement in RF distance is defined as

$$\Delta\text{RF} = \frac{RF(G \setminus \mathcal{O}, S \setminus \mathcal{O}) - RF(G,S)}{1/10 \sum_j RF(G \setminus \mathcal{R}_j, S \setminus \mathcal{R}_j) - RF(G,S)} - 1 = \frac{RF(G \setminus \mathcal{O}, S \setminus \mathcal{O}) - 1/10 \sum_j RF(G \setminus \mathcal{R}_j, S \setminus \mathcal{R}_j)}{1/10 \sum_j RF(G \setminus \mathcal{R}_j, S \setminus \mathcal{R}_j) - RF(G,S)} \tag{4}$$

where $RF$ measures the Robinson and Foulds (1981) distance **without** normalization by the total number of branches. Defining before $= RF(G,S)$, after $= RF(G \setminus \mathcal{R}_j, S \setminus \mathcal{R}_j)$, random $= 1/10 \sum_j RF(G \setminus \mathcal{R}_j, S \setminus \mathcal{R}_j)$, we can write:

$$\Delta\text{RF} = \frac{\text{after} - \text{random}}{\text{random} - \text{before}} \tag{5}$$

Note that since we use RF distance, the denominator of ΔRF score is always positive. Thus, we measure how much more *topologically* similar gene trees become to the species tree due to outlier removal compared to random removal. A positive ΔRF score indicates better than random, and a negative ΔRF score indicates worse than random removal. The logic behind this is that true outliers introduced by error should render gene trees more dissimilar to the species tree than other species.

###### A.2.2 Containment Jaccard

To quantify whether outliers are the same as those found by the branch length-based method TreeShrink, we report the containment Jaccard similarity between the outliers detected by TCMM +PhylteR ($\mathcal{O}_{\text{TCMM+P}}$) and those detected by TreeShrink($\mathcal{O}_{\text{T}}$). Containment Jaccard index is defined as:

$$\text{ContainmentJaccard}(\mathcal{O}_{\text{TCMM+P}}, \mathcal{O}_{\text{T}}) = \frac{(\mathcal{O}_{\text{TCMM+P}} \cap \mathcal{O}_{\text{T}})}{|\mathcal{O}_{\text{TCMM+P}}|}\,.$$

##### A.3 Outlier removal for branch length estimation

There are two types of outliers that need to be considered and we deal with them separately since they have very different properties.

**Small outliers.** TCMM tends to assign exceedingly small values to *some* branches that are in the query but not the reference tree (as we will see), creating outliers. To detect and remove these outliers, we apply Jenks natural breaks to the log-transformed branch length estimates. The log transformation helps by converting the skewed branch length distribution into a more bimodal one, where the outliers form a distinct group of extremely small values easily separable by a clustering method such as Jenks natural breaks. We identify a single breakpoint and remove the data below the breakpoint only if splitting the data at this point reduces the variance by more than fivefold. The variance reduction is calculated for the two equal-sized sets of data points, $x1$ and $x2$, using the formula:

$$2\frac{\sigma^2(x_1\|x_2)}{(\sigma^2(x_1)+\sigma^2(x_2))}$$

where $x_1\|x_2$ denotes the concatenation of the two sets. Since the two resulting groups may differ in size, we balance them by randomly sampling from the larger group to match the size of the smaller one.

**Large outliers.** The per-gene method can also introduce extremely high values, which can arise from aberrantly long branches in the the gene trees or specific forms of discordance. These kinds of outliers are much less common than small values. As a result, they cannot be effectively separated using the same method as for lower outliers. To address these extreme high values, we use the simple boxplot method: we eliminate any data point exceeding $Q_3 + 3 \times IQR$, where $Q_3$ is the third quartile and $IQR = Q_3 - Q_1$ is the interquartile range. Note that this step follows the previous step, ensuring the lower values are removed first. This approach helps minimize the impact of outliers on the final branch length estimates, improving accuracy (as shown in the results).

#### A.4 Missing data

For a branch $e \in E_T$, let $p_T(e)$ represent the number of leaf pairs $(u, v)$ where $u, v \in L_T$ such that $e \in P_T(u, v)$, and let $p^*(e)$ be the number of such leaf pairs that also exist in $R$; note $p^*(e) \leq p_T(e)$, with equality indicating no missing data. When $p^*(e) < p_T(e)$, the value of $(A_T^\top d_R)_e$ tends to be underestimated. To see this, consider a case where $R$ and $T$ are identical and note that $(A_T^\top d_R)_e$ gives the sum of patristic distances that include $e$ on their path. Now if some leaves are removed in $R$, that sum necessarily decreases. We propose two solutions to reduce this bias. The first approach simply scales $A_T^\top d_R$ by multiplying each $(A_T^\top d_R)_e$ with $p_T(e)/p^*(e)$ if $p^*(e) > 0$. This approach assumes that the pairs of taxa missing from $R$ have similar mean patristic distances to those that are present. While we can construct cases where this assumption is violated (e.g., when outgroups are missing), this assumption seems reasonable empirically (See Fig. S2), and the method largely eliminates the underestimation bias.

The scaling method, however, fails when $p^*(e) = 0$, meaning that no pair around an edge is present in $R$. An alternative approach that works even when $p^*(e) = 0$ is imputation using neighbors. In this method, for each leaf $l \in L_T \setminus L_R$, the closest leaf $l' \in L_R$ is selected and $l$ is added as a sibling of $l'$ to $R$ to form a cherry with both terminal branches set to zero.

We deal with the missing data slightly differently depending on whether consensus or per-gene modes are used. In the consensus version (including all cases of application 1), for each branch $e$, we apply the scaling heuristic mentioned earlier for gene trees that include at least one pair around $e$ and ignore gene trees that do not have any; we assume that around every edge $e$, there is at least one pair that appears together in at least one gene tree.

To deal with missing data in the per-gene mode, we use a combination of scaling and placement heuristics. For any branch $e$ in the species tree, if none of the pairs around it appear in a particular gene tree, we use the placement method to add it; otherwise, we apply the scaling heuristic to fix $A_T d$ values for that gene tree.

#### A.5 Supplementary remarks and proofs

**Remark 1.** *Problem 3 reduces to Problem 2.*

*Proof.* We can rewrite the objective function of Problem 3 as

$$\sum_{k\in[K]} \alpha_k \|d_{R_k} - A_T w\|_2^2 + \lambda\sigma^2(w/w_T) = w^\top A_T^\top A_T w\big(\sum_{k\in[K]} \alpha_k\big) - 2\big(\sum_{k\in[K]} \alpha_k d_{R_k}\big)^\top A_T w + \lambda\sigma^2(w/w_T) + \text{const}$$

$$= \text{const}''\|\sum_{k\in[K]} \frac{\alpha_k}{\sum_{n\in[K]}\alpha_n} d_{R_k} - A_T w\|_2^2 + \lambda\sigma^2(w/w_T) + \text{const}' ,$$

which equals Eq. (2) if we set

$$d = \sum_{k\in[K]} \frac{\alpha_k}{\sum_{n\in[K]}\alpha_n} d_{R_k} .$$

Thus, we simply need to compute the weighted average of reference distance vectors and solve the problem with this matrix as input. □

**Remark 2** (Equivalency of Consensus and Per-gene average)**.** *Consider the consensus Problem 3 with uniform weights ($\alpha_k = 1$ for all $k \in [K]$) and Problem 2 applied to each reference tree separately. Although both problems are constrained by $w \geq \varepsilon$, the unconstrained global minimizer of their objective may lie outside this region (i.e., some entries of $w^*$ may be less than $\varepsilon$). Consider only cases when all optimal unconstrained solutions (branch vectors) do lie strictly within the feasible region, i.e., are greater than or equal to $\varepsilon$. In these cases, the arithmetic average of solutions to Problem 2 is also the solution to the consensus Problem 3.*

*Proof.* Let us assume that the unconstrained optimal branch vector $w^*$ lies within the feasible region, i.e.,

$w^* \geq \varepsilon$. Then, $w^*$ satisfies the first-order optimality condition:

$$\nabla_w \frac{1}{2K} \sum_{k\in[K]} \|d_{R_k} - A_T w\|_2^2 + \lambda\sigma^2(w/w_T)\Big|_{w=w^*} = A_T^\top A_T w^* - A_T^\top \left(\frac{1}{K}\sum_{k\in[K]} d_{R_k}\right) + \lambda\frac{2}{|E_T|}P_1^\perp(w^*/w_T^2) = 0.$$

Let us now compute the optimal solution to the regularized version of Problem 2 between $T$ and each $R_k$:

$$\forall k \in [K] : \nabla_w \frac{1}{2}\|d_{R_k} - A_T w\|_2^2 + \lambda\sigma^2(w/w_T)\Big|_{w=w_k^*} = A_T^\top A_T w_k^* - A_T^\top d_{R_k} + \lambda\frac{2}{|E_T|}P_1^\perp(w_k^*/w_T^2) = 0.$$

This holds statement *because* we assumed that the unconstrained optimal branch vectors $w_k^*$ for all references lie within the feasible region ($w_k^* \geq \varepsilon$). If the optimal unconstrained solution was outside the region, we could not have asserted that the gradients are zero at the optimal solution. Averaging both sides of this equation across $k \in [K]$, we obtain

$$A_T^\top A_T \left(\frac{1}{K}\sum_{k\in[K]} w_k^*\right) - A_T^\top \left(\frac{1}{K}\sum_{k\in[K]} d_{R_k}\right) + \lambda\frac{2}{|E_T|}P_1^\perp\big((\frac{1}{K}\sum_{k\in[K]} w_k^*)/w_T^2\big) = 0\ ,$$

which matches the condition for the consensus problem, that is, $w^* = \frac{1}{K}\sum_{k\in[K]} w_k^*$. □

*Proof of Proposition 1.* The gradient of the cost function in Eq. (3) with respect to the weights $w \in \mathbb{R}^{|E|}$ is:

$$\nabla_w l_1(w) = Bw + c,$$

where $B \in \mathbb{R}^{|E|\times|E|}$ and $c \in \mathbb{R}^{|E|}$. The Hessian $\nabla_w^2 l_1(w) = B$ is a positive semidefinite matrix; thus, $l_1(w)$ is a convex function everywhere in its domain. We can also compute the gradient of $l_2(w)$ as follows:

$$\nabla_w l_2(w) = \frac{2}{|E|}(P_1^\perp r)(1/w_T) = \frac{2}{|E|}P_1^\perp(w/w_T^2).$$

Furthermore, its Hessian $\nabla_w^2 l_2(w) = \frac{2}{|E|}\mathrm{diag}(1/w_T)P_1^\perp\mathrm{diag}(1/w_T)$ is a positive semidefinite matrix, making $l_2(w)$ a convex function of $w$ as well. Since both $l_1(w)$ and $l_2(w)$ are convex functions of weight parameter $w$ and cost functions in Problems 1, 2, and 3 are linear functions of $l_1$ and $l_2$ with *positive coefficients* ($\lambda$ and $\alpha_k^2$ terms), each cost function is also a convex function of $w$. In Problem 1, the constraint set $w \geq 0$ is nonempty, closed, and convex. In Problems 2 and 3, the constraint set $w \geq \varepsilon$ is also nonempty, closed, and convex. Therefore, all three problems involve convex function minimizations over nonempty, closed, and convex sets. Since convex functions over nonempty, bounded, convex sets have a minimum, the projected gradient descent method converges to the global minimum of the cost functions (Wang and Xiu, 2000). □

**Algorithm S1** Computing $A_T^\top A_T$ and $A_T^\top d_R$

**Notations:** $e(n)$ is the edge above $n$. $C_T(e)$ is the set of leaves in $T$ under edge $e$.

```
procedure COMPUTE_$A^\top A$(Tree $T$)
    pre-compute $|C_T|$ for each edge of $T$ in a bottom-up traversal
    for $e$ in $E_T$ do
        for $e'$ in $E_T$ do
            if $e == e'$ then
                $A^\top A[e,e'] \leftarrow |C_T(e)| \cdot (|L_T| - |C_T(e)|)$
            else
                if $e$ is a descendant of $e'$ then
                    $A^\top A[e,e'] \leftarrow |C_T(e)| \cdot (|L_T| - |C_T(e')|)$
                else if $e'$ is a descendant of $e$ then
                    $A^\top A[e,e'] \leftarrow |C_T(e')| \cdot (|L_T| - |C_T(e)|)$
                else
                    $A^\top A[e,e'] \leftarrow |C_T(e)| \cdot |C_T(e')|$
    return $A^\top A$

procedure TOTAL_PDISTANCE_TO_LEAF(Leaf $l$, $C'$, $L'$, Tree $T'$)    ▷ Computes sum of distances to $l$ from all leaves
    $distance \leftarrow 0$
    $length \leftarrow 0$
    $node \leftarrow l$
    while $node \neq \text{root}(T')$ do
        $m \leftarrow$ sibling of $node$
        $length$ += $w_{e(node)}$
        $distance$ += $L'(e(m)) + length \cdot |C'(e(m))|$
        $node \leftarrow \text{parent}(node)$
    return $distance$

procedure COMPUTE_$A^\top d$(Tree $T$, Tree $R$)
    Bottom-up traverse $R$ to compute $C_R(e')$ and $L_R(e') = \sum_{w \in C_R(e')} d_R(w,u')$ for all $e' = (u',v') \in E_R$
    Bottom-up traverse $T$ to compute $C_T(e)$ for $e \in E_T$
    for $n \in$ bottom-up traversal of $V_T$ do
        if $n$ is leaf then
            $n' \leftarrow n$ in $R$
            $A^\top d[e(n)] = \text{total\_pdistance\_to\_leaf}(n', C_R, L_R, R)$
        else
            $c_l, c_r \leftarrow$ children of $n$
            $A^\top d[e(n)] \leftarrow A^\top d[e(c_l)] + A^\top d[e(c_r)]$
            $C_l(e), L_l(e) \leftarrow \emptyset, 0$ for $e \in E_T$
            for $n_l \in C_T(e(c_l))$ do
                $n'_l \leftarrow n_l$ in $R$
                $C_l(e(n'_l)) \leftarrow \{n'_l\}$
                $L_l(e(n'_l)) \leftarrow w_{e(n'_l)}$
                while $n'_l \neq \text{root}(R)$ do
                    $c'_1, c'_2 \leftarrow$ children of $n'_l$
                    $C_l(e(n'_l)) \leftarrow C_l(e(c'_1)) \cup C_l(e(c'_2))$
                    $L_l(e(n'_l)) \leftarrow L_l(e(c'_1)) + L_l(e(c'_2)) + |C_l(e(n'_l))| \cdot w_{e(n'_l)}$
                    $n'_l \leftarrow \text{parent}(n'_l)$
            for $n_r \in C_T(e(c_r))$ do
                $n'_r \leftarrow n_r$ in $R$
                $A^\top d[e(n)]$ −= $2 \cdot \text{total\_pdistance\_to\_leaf}(n'_r, C_l, L_l, R)$
    return $A^\top d$
```

### B Supplementary Tables

Table S1: Simphy parameters used in S100 simulations. The parameters not mentioned in the second and third tables follow the default parameters of Zhang et al. (2018).

| **Parameter Name** | **Parameter Value** |
|---|---|
| Default S101 data by by Zhang et al. 2018 for the S100 dataset | |
| Speciation rate (`sb`) | 0.0000001 |
| Extinction rate (`sd`) | 0 |
| Number of Leaves | 100 |
| Ingroup to outgroup ratio (`so`) | 1.0 |
| Generations (`st`) | LogN(1.470055e+01,2.500000e-01) |
| Haploid effective population size (`sp`) | 400000 |
| Global substitution rate (`su`) | LogN(-1.727461e+01,6.931472e-01) |
| Species-specific rate gamma shape (`hs`) | LogN(1.5e+00,1) |
| Gene-specific rate gamma shape (`hl`) | LogN(1.551533e+00,6.931472e-01) |
| Gene-by-lineage rate gamma shape (`hg`) | LogN(1.5e+00,1) |
| Seed | 9644 |
| Sequence Length | 1600, 800, 400, 200 |
| Sequence base frequencies | Dirichlet(A=36,C=26,G=28,T=32) |
| Sequence transition rates | Dirichlet(TC=16,TA=3,TG=5,CA=5,CG=6,AG=15) |
| S100-truegt - varying ILS | |
| Haploid effective population size (`sp`) | $4 \times 10^4$: low, $4 \times 10^5$: medium, or $4 \times 10^6$: high ILS |
| Lineage specific rate gamma shape (`hs`) | NA (i.e., no rate heterogenity across species) |
| S100-truegt - varying rate heterogenity | |
| Species-specific rate gamma shape (`hs`) | LogN(1.5e+00,1) in `hl+hs` and `hl+hg+hs`; NA in `hl+hg` |
| Gene-by-lineage rate gamma shape (`hg`) | LogN(1.5e+00,1) in `hl+hg` and `hl+hg+hs`; NA in `hl+hs` |

Command used to generate:

```
# Default S100 dataset (from original paper)
./simphysu -rs 50 -rl f:1000 -rg 1 -sb f:0.0000001 -sd f:0 -st ln:14.70055,0.25
-sl f:100 -so f:1 -si f:1 -sp f:400000 -su ln:-17.27461,0.6931472 -hh f:1 -hs ln:1.5,1
-hl ln:1.551533,0.6931472 -hg ln:1.5,1 -cs 9644 -v 3 -o ASTRALIII -ot 0 -op 1 -od 1

# S100 lowILS:
    Change -sp f:400000 --> -sp f:40000
    Remove -hs ln:1.5,1

# S100 mediumILS:
    Remove -hs ln:1.5,1

# S100 highILS:
    Change -sp f:400000 --> -sp f:4000000
    Remove -hs ln:1.5,1

# S100 hl+hs:
    Remove -hg ln:1.5,1

# S100 hl+hg:
    Remove -hs ln:1.5,1

# S100 hl+hg+hs:
    No changes.
```

### C Supplementary figures

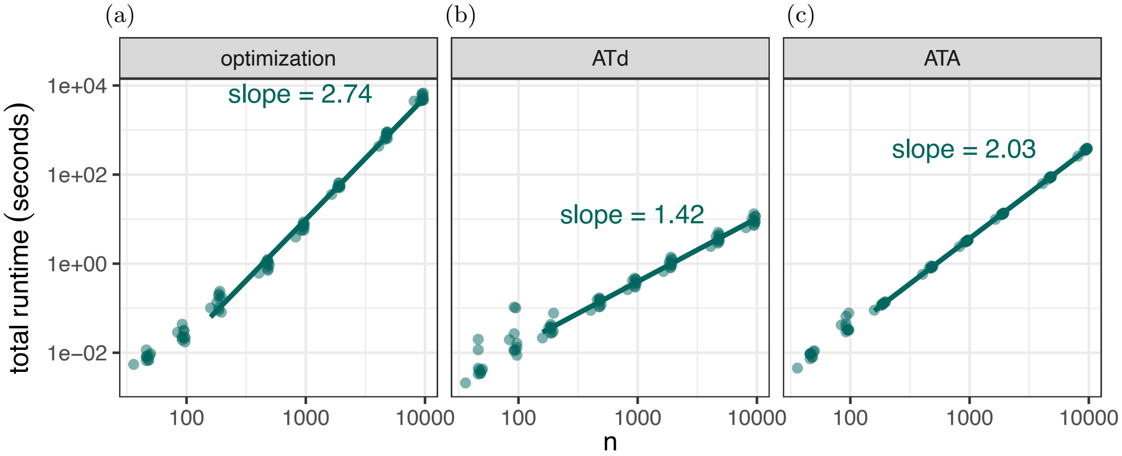

Figure S1: Running time of the (a) optimization, (b) computing $A_T^\top d_R$, and (c) computing $A_T^\top A_T$ step of TCMM with respect to the size of $T$. Starting from the real dataset of Zhu et al. (2019) with 10575 leaves, for each $n \in \{50, 100, 200, 500, 1000, 2000, 5000, 10000\}$, we prune the species tree $T$ and the gene tree $R$ to contain only $n$ taxa. If some taxa are absent from the gene tree, we remove them from the species tree so that both $T$ and $R$ are the same size. The slopes in the log-log plots show the empirical complexity of each step of TCMM. Each point represents running TCMM on the pruned species tree and one pruned gene tree. We used the first ten gene trees in this dataset and fit the line to the mean across the 10 runs, excluding the small $n \leq 100$ in line fitting.

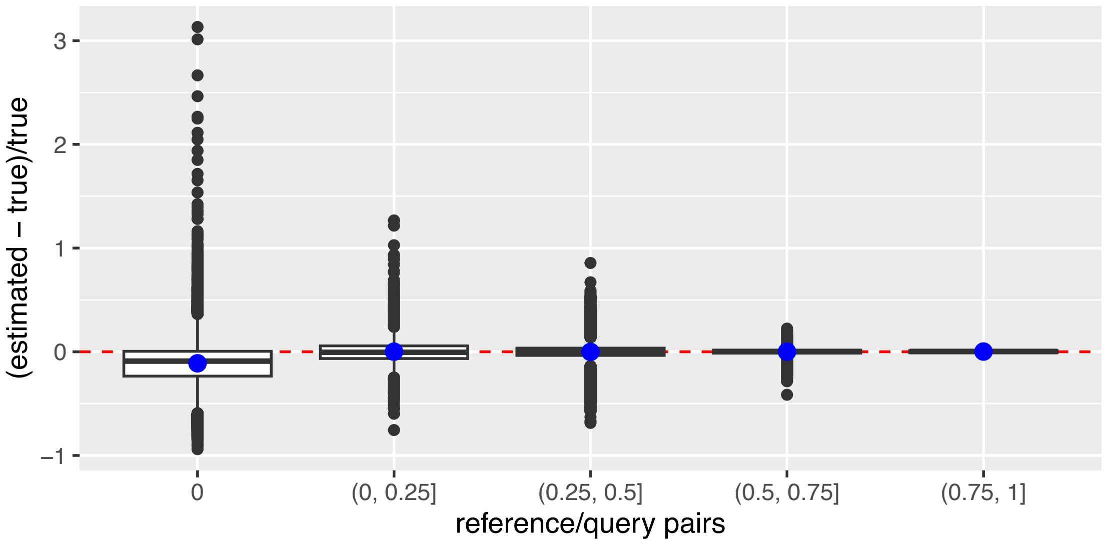

Figure S2: (S100-default Dataset, true gene trees) Comparing estimated values for $A_T^\top d_R$ to the true values. Missing taxa are introduced into the true gene trees by removing $m_g$ species from gene tree $g$, where $m_g \sim \text{Poisson}(\lambda = 0.34n)$ and $n$ is the total number of leaves in the complete gene trees. The scaling method is used to estimate $(A_T^\top d_R)_e$ where $p_*(e) > 0$ for a branch $e$ and the imputation method is used otherwise. The true values are computed using complete true gene trees. The figure shows results for the first gene tree as the query and the first 10 gene trees as the reference trees (pruned trees for estimated values and complete trees for true values).

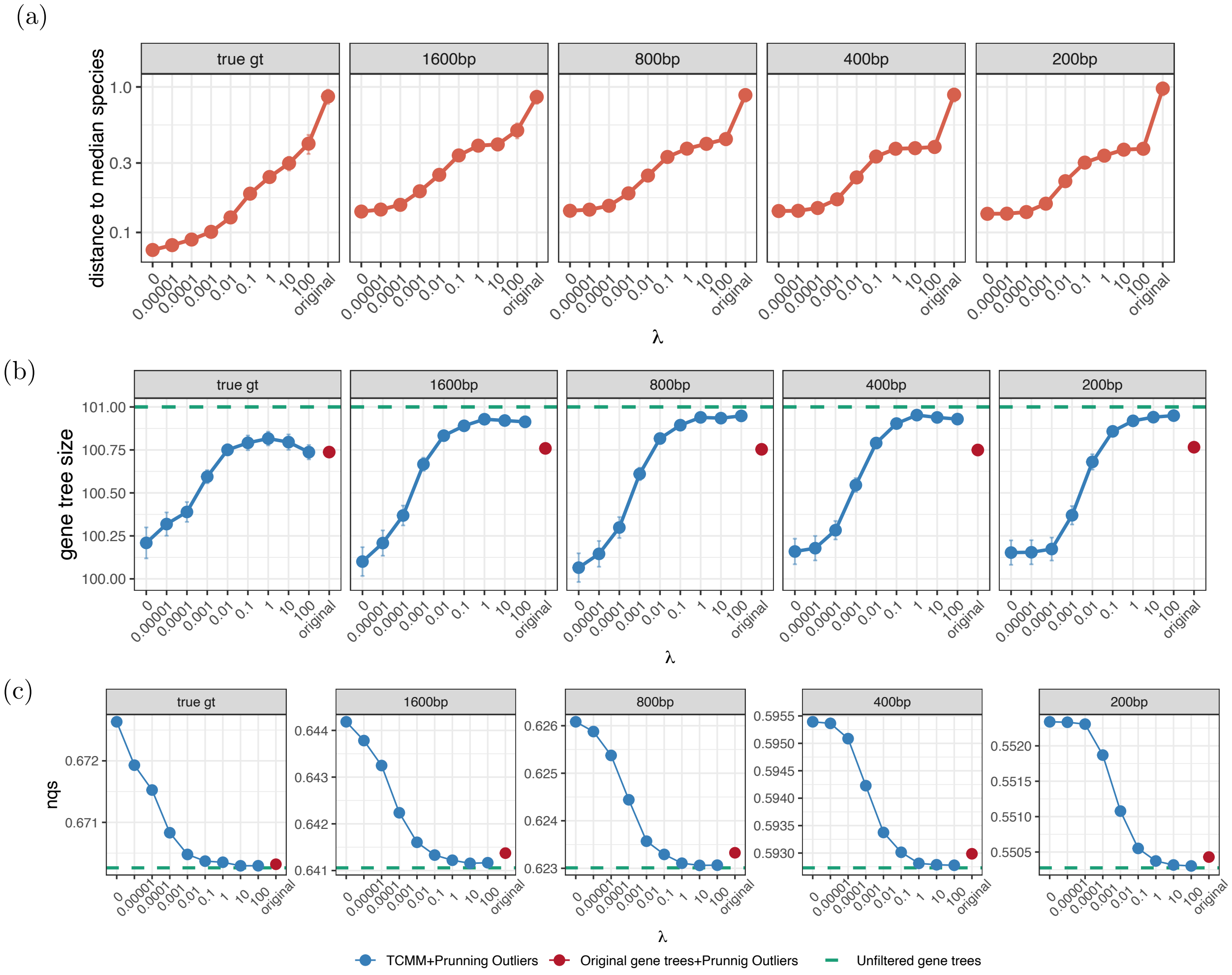

Figure S3: (S100-default Dataset) (a) Comparing the Euclidean distance to the median species for all levels of gene tree error. (b) Size of gene trees normalized by total number of species per replicate, averaged across all replicates. Mean and standard error are shown for each $\lambda$. (c) Normalized quartet score to the true species tree for different levels of gene tree error. The unfiltered estimated and true gene trees are shown in green.

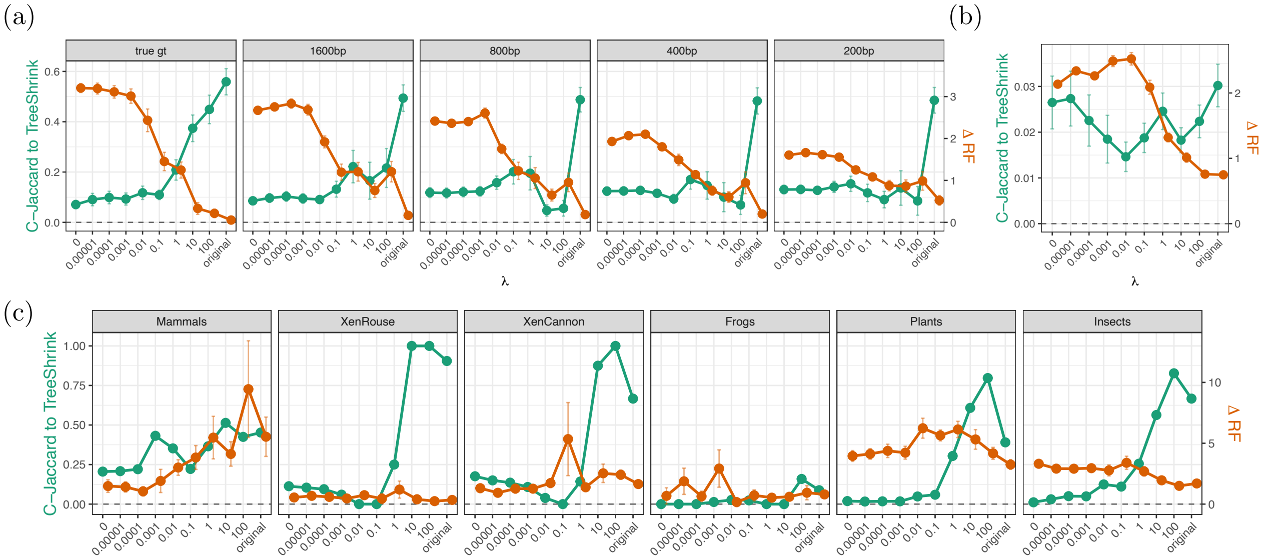

Figure S4: Similarity to TreeShrink outliers in Containment Jaccard (green) and ΔRF distance (orange) for (a) S100-default, (b) S200-perturbed, and (c) biological datasets. ΔRF is defined as the decrease in the RF distance of gene trees to the species tree after removing outliers, normalized by the same quantity with random removal of the same number of leaves (Appendix A.1). Above 0 indicates better than random.

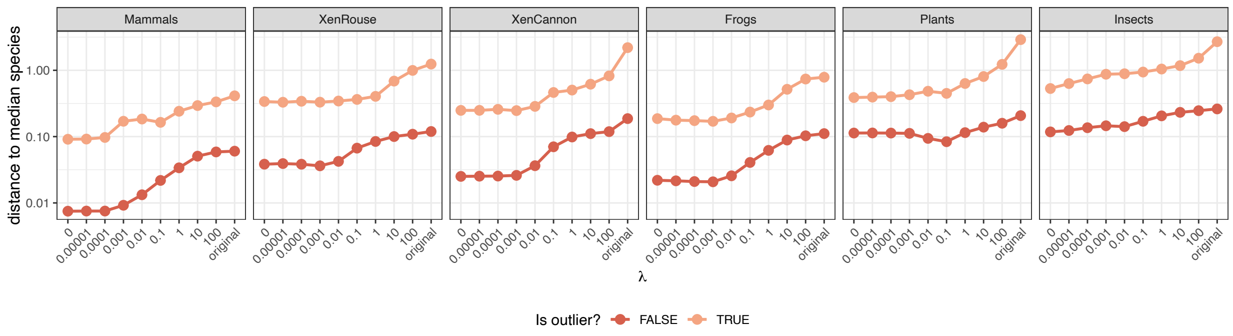

Figure S5: (Biological Dataset) Comparing the Euclidean distance to the median species for outlier and non-outlier species.

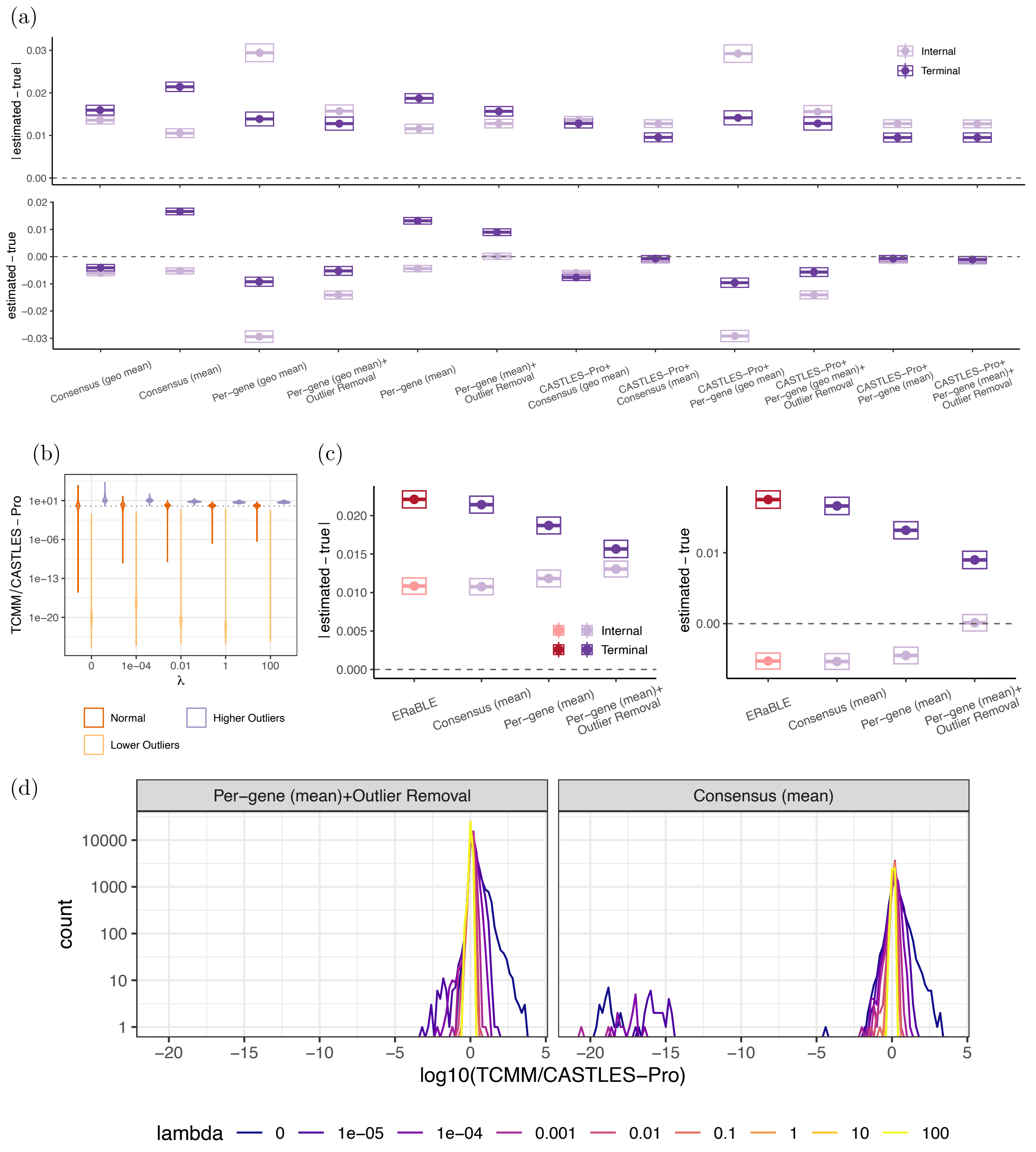

Figure S6: (HGT Dataset, HGT rate = 2e-7) (a) Comparison of different versions of TCMM in terms of mean absolute error (top) and bias (bottom), showing mean (dot) and 0.95% normal CI (box). Summarization methods are arithmetic mean (*mean*) or geometric mean (*geo mean*). The regularization coefficient ($\lambda$) was set to zero, except for the CASTLES-Pro+per-gene and CASTLES-Pro+consensus method, where $\lambda$ is selected automatically. (b) The distribution of the estimated branch lengths of two versions of TCMM divided by the CASTLES-Pro branch lengths. (c) Comparison of different versions of TCMM with the distance-based method ERaBLE in terms of mean absolute error (left) and bias (right), showing mean (dot) and 0.95% normal CI (box). (d) The effect of the regularization coefficient ($\lambda$) on the estimated branch lengths. TCMM output length per gene (before summarization) over the input length (CASTLES-Pro) is shown; the variance of this ratio (spread of distribution) indicates the change in the relative length of branches between input and output of TCMM. Outliers are identified by our per-gene method.

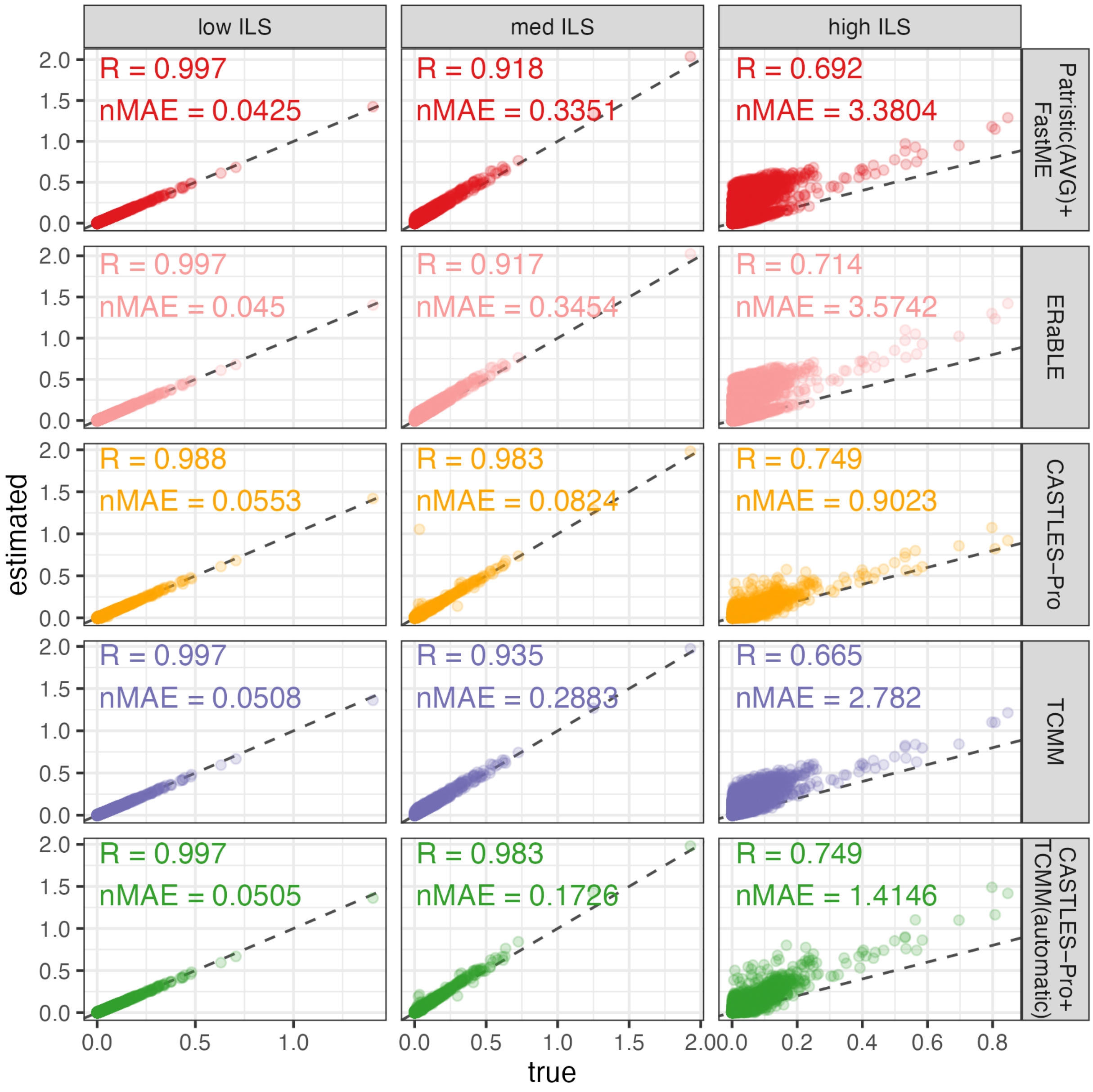

Figure S7: (S100-truegt Dataset, hl+hg) The scatter plot of true branch lengths vs. estimated branch lengths for different levels of ILS. Spearman correlation (R) and mean absolute error normalized by mean true branch lengths (nMAE) are shown for each method.

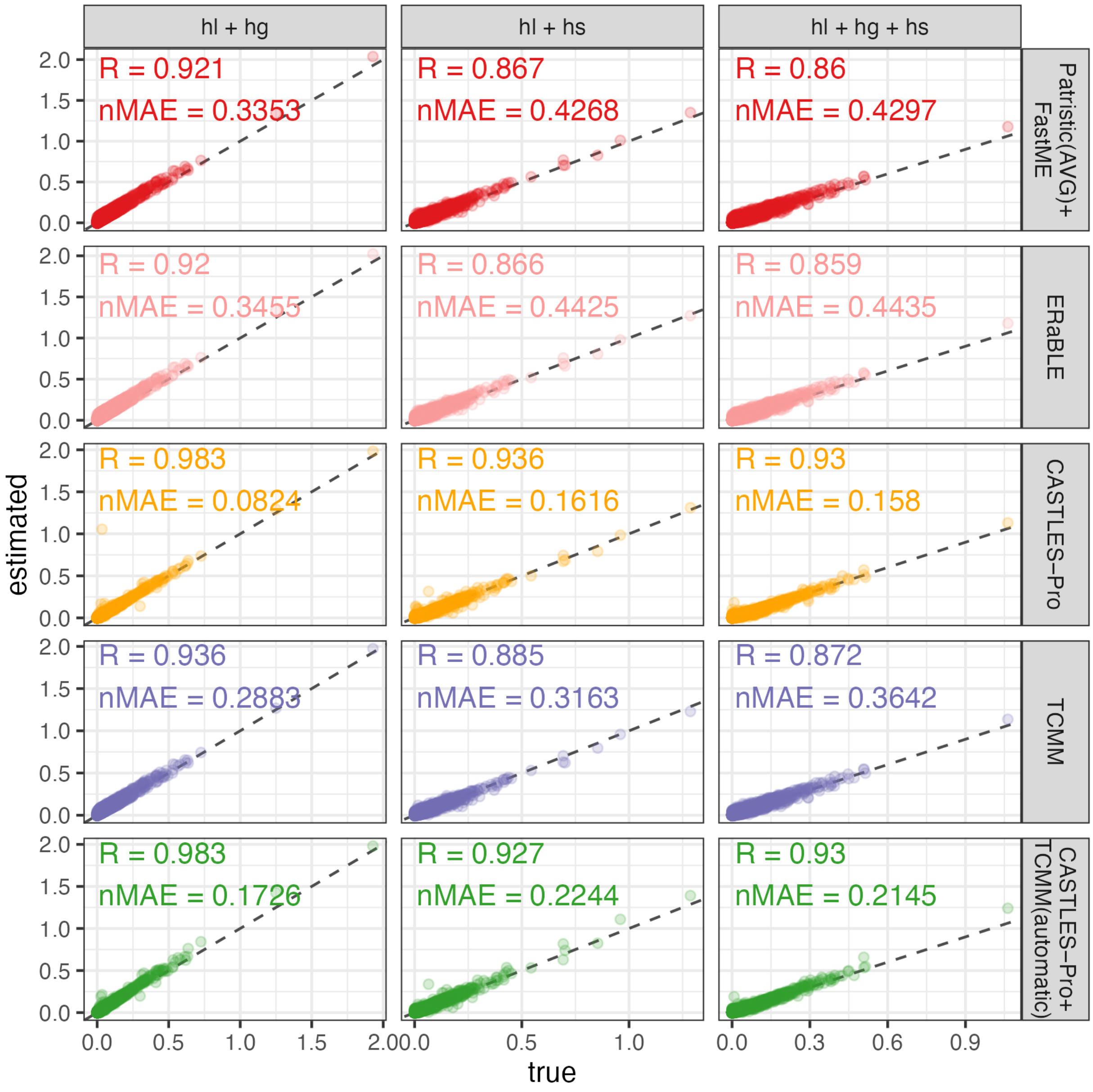

Figure S8: (S100-truegt Dataset, medium ILS) The scatter plot of true branch lengths vs. estimated branch lengths for different levels and types of heterogeneity. Spearman correlation (R) and mean absolute error normalized by mean true branch lengths (nMAE) are shown for each method.

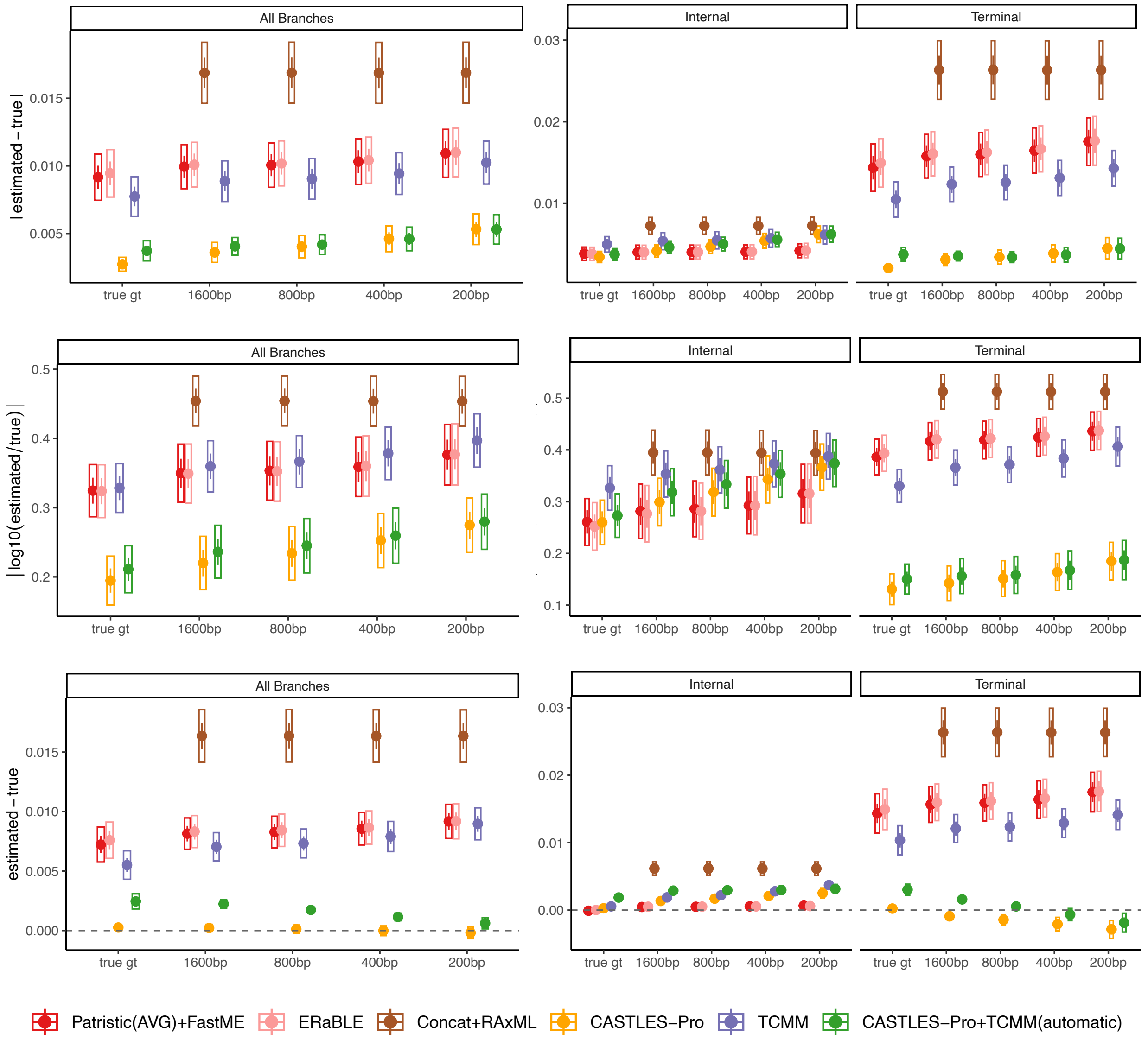

Figure S9: (S100-default Dataset) Comparing TCMM with the alternative methods across various levels of gene tree error (x-axis). TCMM is run with $\lambda = 0$ (TCMM) and automatic $\lambda$ selection (CASTLES-Pro+TCMM). The methods are compared using two error metrics (mean absolute error and mean absolute log error) as well as bias.

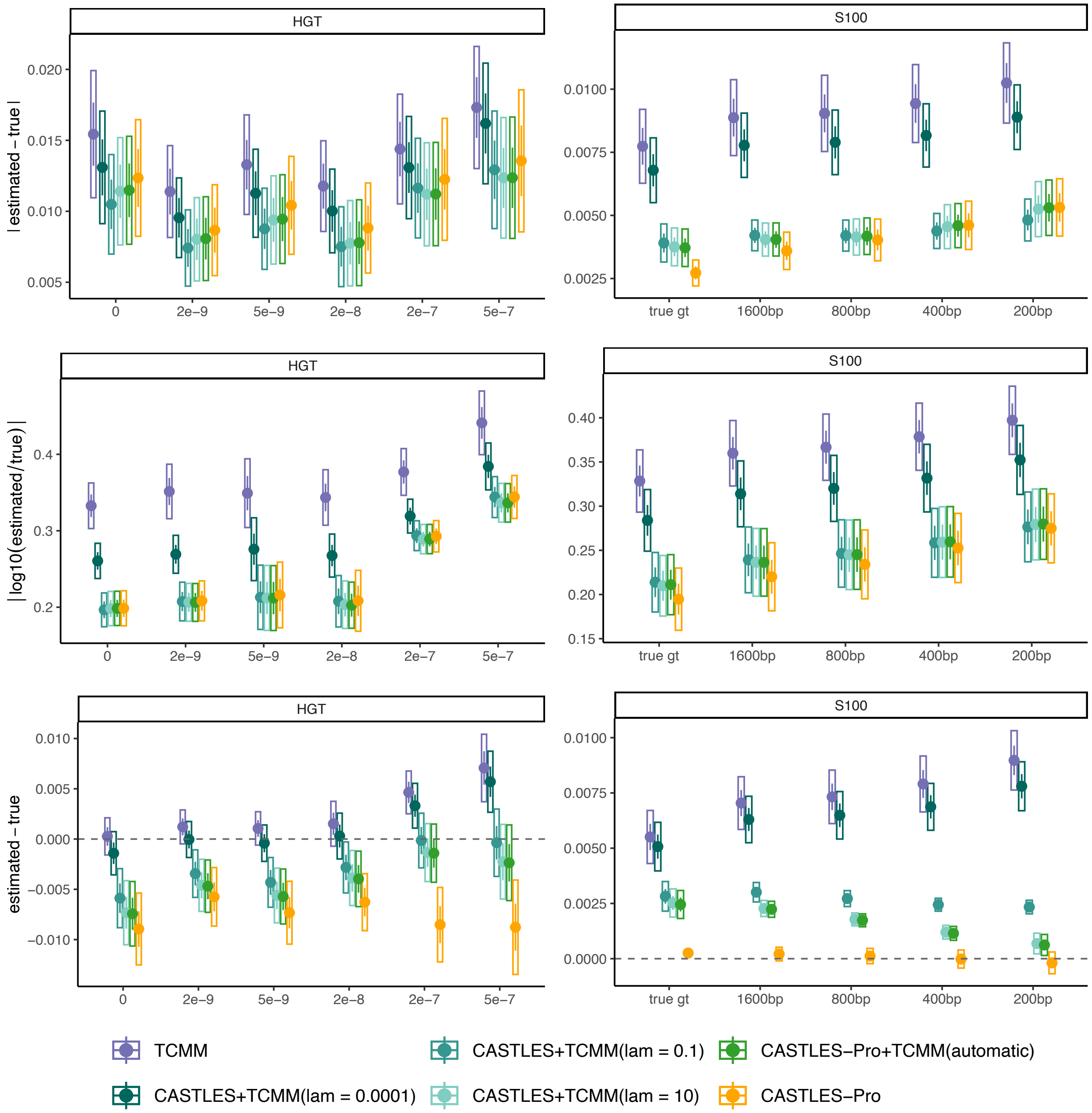

Figure S10: Comparing different $\lambda$ values for CASLTES-Pro+TCMM (including automatic $\lambda$ selection) with CASTLES-Pro and TCMM alone on the HGT dataset (left) and S100-default dataset (right). The methods are compared using two error metrics (mean absolute error and mean absolute log error) as well as bias.

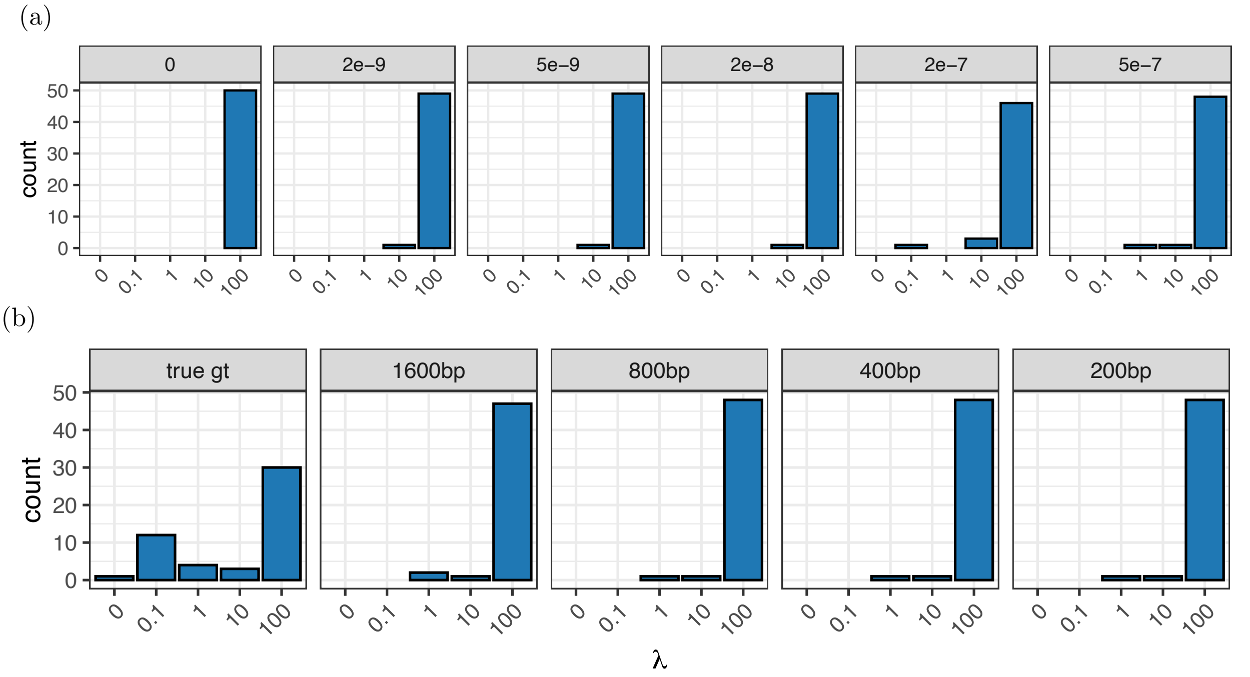

Figure S11: Histogram of the $\lambda$ chosen by automatic $\lambda$ selection process of TCMM for different model conditions for the HGT dataset (a) and S100-default dataset (b).
